# Spatial organization and stochastic fluctuations of immune cells impact clinical responsiveness to immune checkpoint inhibitors in patients with melanoma

**DOI:** 10.1101/2023.12.06.570410

**Authors:** Giuseppe Giuliani, William Stewart, Zihai Li, Ciriyam Jayaprakash, Jayajit Das

## Abstract

High-dimensional, spatial single-cell technologies such as CyTOF imaging mass cytometry (IMC) provide detailed information regarding locations of a large variety of cancer and immune cells in microscopic scales in tumor microarray (TMA) slides obtained from patients prior to immune checkpoint inhibitor (ICI) therapy. An important question is how the initial spatial organization of these cells in the tumor microenvironment (TME) change with time, regulate tumor growth and eventually outcomes as patients undergo ICI therapy. Utilizing IMC data of melanomas of patients who later underwent ICI therapy, we develop a spatially resolved interacting cell systems model that is calibrated against patient response data to address the above question. We find that the tumor fate in these patients is determined by the spatial organization of activated CD8+ T cells, macrophages, and melanoma cells and the interplay between these cells that regulate exhaustion of CD8+ T cells. We find that fencing of tumor cell boundaries by exhausted CD8+T cells is dynamically generated from the initial conditions that can play a pro-tumor role. Furthermore, we find that specific spatial features such as co-clustering of activated CD8+ T cells and macrophages in the pre-treatment samples determine the fate of the tumor progression, despite stochastic fluctuations and changes over the treatment course. Our framework enables determination of mechanisms of interplay between a key subset of tumor and immune cells in the TME that regulate clinical response to ICIs.

**Significance:** Recent advances in single cell technologies allows for spatial imaging a wide variety of cancer and immune cells in tissue samples obtained from solid tumors. This detailed snapshot data of microscale organization of tumor and immune cells could provide valuable insights into underlying biology and clinical responsiveness to cancer immunotherapy. By combining published data from imaging mass-cytometry and patient response against ICI drugs with data analysis rooted in statistical physics and statistical inference theory, we developed and studied the dynamics of mechanistic spatially resolved models: we show that tumor growth during ICI treatment is regulated by non-intuitive interplay between CD8+ T cells and tumor associated macrophages, formation of a pro-tumor fencing of exhausted CD8+ T cells around melanoma cells, specific features of spatial organization of these cells prior to treatment, and stochastic fluctuations in the dynamics. The mechanisms unveiled in our studies are general and can pertain to the response of other solid tumors to ICI therapy.

## Introduction

A tumor is complex and heterogeneous tissue composed of tumor cells, various immune cells, connective tissues, blood and lymphatic vessels, and extracellular materials like collagen (2). The relationship between tumor cells and immune cells in the tumor microenvironment (TME) is shaped by multiple factors, including the recruitment of immune cells (3), activation of T cells by neoantigens in the draining lymph nodes, and exhaustion of activated T cells. Immunotherapeutic strategies that exploit the immune system to eliminate tumors have revolutionized cancer treatment (4). However, despite many striking examples of success these therapies still cure only a moderate percentage (∼20-40%) of patients (4, 5). Systems level understanding of how the immune system induces anti- and pro-tumor responses can considerably aid the development of newer immunotherapies with greater efficacy.

Recent advancements in single cell spatial technologies such as CyTOF imaging mass cytometry (IMC)(6, 7) and Cyclic Immunofluorescence (CyCIF)(1, 8) provide extensive information about the composition of the TME where over thirty different proteins can be measured in single cells along with knowledge of their spatial locations in small millimeter scale tissue microarray (TMA) samples extracted from tumors. A wide variety of spatial data analysis tools ranging from calculation of local cell density (9–11), spatial pair correlations (10, 12), clustering (13) to application of probabilistic methods such as latent Dirichlet allocation (12, 14) and machine learning methods such as graph-(15) or convolutional-(16) neural networks have been employed to identify microscopic spatial patterns of tumor and immune cells in high dimensional IMC, CyCIF, and multispectral fluorescence imaging datasets that are associated with patient survival and response to anti-cancer drugs. However, the imaging is usually done at a single timepoint, mostly before the ICI treatment, and a major challenge is to infer how these microscopic spatial patterns mechanistically determine the tumor growth kinetics when checkpoint drugs are administered.

Spatially resolved mechanistic models involving tumor and immune cells have been developed over the years using agent based(17–21) and cellular automaton(20) modeling approaches or partial differential equations(22) to determine the roles of interaction between these cell types, vascularization, and mechanical forces in tumor growth and patient responses. These models largely remained uncalibrated against experimental data, however, due to the access of digital H&E slides in the recent years, greater computation power, and efficient algorithms, several studies have calibrated spatial models against real(23) and synthetic(24) immunohistochemistry datasets which can identify few types of cells. In contrast, IMC datasets can identify many immune cell types as well their activation states which offer an attractive scenario to calibrate such spatial models and infer mechanisms involving interplay between the tumor and immune cells.

To this end, we combined IMC data obtained from biopsy samples from melanoma patients pre-checkpoint therapy, the data regarding the response of these patients to anti-PD1/CTLA4 checkpoint therapy to develop a mechanistic spatially resolved interacting cell systems model; we combined statistical interference theory to justify details of the interactions between immune and tumor cells that underlie patient responses to immune checkpoint inhibition (ICI) drugs. The specific major findings from our investigation are the following: (i) The elucidation of the effects of the interplay between killing of tumor cells by active CD8+ T cells, and the exhaustion of the latter by tumor cells and non-inflammatory macrophages, (ii) how the interplay in (i) dynamically generates spatial configurations with fencing of tumor cells by exhausted CD8+ T cells protecting tumor cells from active CD8+ T cells outside the fence; and (iii) how, despite the inherent stochastic fluctuations in the processes, at time scales longer than cell proliferation, maturation and death, specific features in the initial spatial organization of the melanoma and immune cells lead to widely different tumor growth outcomes. Our modeling approach and methods can be utilized to design new immunotherapeutic strategies for melanoma and other cancers.

### Approach

We obtained publicly available CyTOF imaging mass cytometry (IMC) data of 1×1 mm^2^ tissue microarrays (TMA) to quantify proteins in single cells that yield a spatial profile of different cell types in the tumor microenvironment (TME). This information for samples from a cohort of 30 melanoma patients is paired with the response (responder or non-responder) to immune checkpoint inhibitors (ICIs) involving anti-PD1, anti-CTLA4, or the combination of the two (6).

We followed three main steps. First, we used the IMC and the patient response datasets to identify the immune cell types and their microscale spatial organization that are associated with the response of patients to ICI drugs. Second, we developed a spatially resolved interacting cell system (ICS) model involving mechanistic interactions between tumor and the relevant immune cell types identified in the previous step. Lastly, we set up a training and testing framework for the spatially resolved mechanistic model using the IMC and the patient response datasets; this allows us to evaluate different hypotheses regarding the interactions underlying the interplay between tumor and immune cells. We describe these steps briefly below and provide further details in the Materials and Methods section and in the Supplementary Material.

#### Identification of relevant spatial microscale cellular patterns

We used the spatial data regarding the locations of 10 different cell types such as melanoma cells, activated CD8+ T cells, and macrophage/monocytes in the TMA slides that were determined by (6) for our analysis. First, we computed the densities of these different cell types in each TMA slides which displayed large patient-patient variations. To determine cell types whose densities differ substantially between responders (14 patients) and non-responders (16 patients) of the ICI therapy, we compared the mean densities of the cell types averaged over TMA slides obtained from responder or non-responder patients (Table S1a). We found that responders have on average larger densities of activated CD8+ T cells (p=0.16) (Fig. 1a); however, the densities of all the cell types did not display substantial difference (p≤ 0.05) between the responder and non-responders (Fig. S1a).

**Figure 1.**
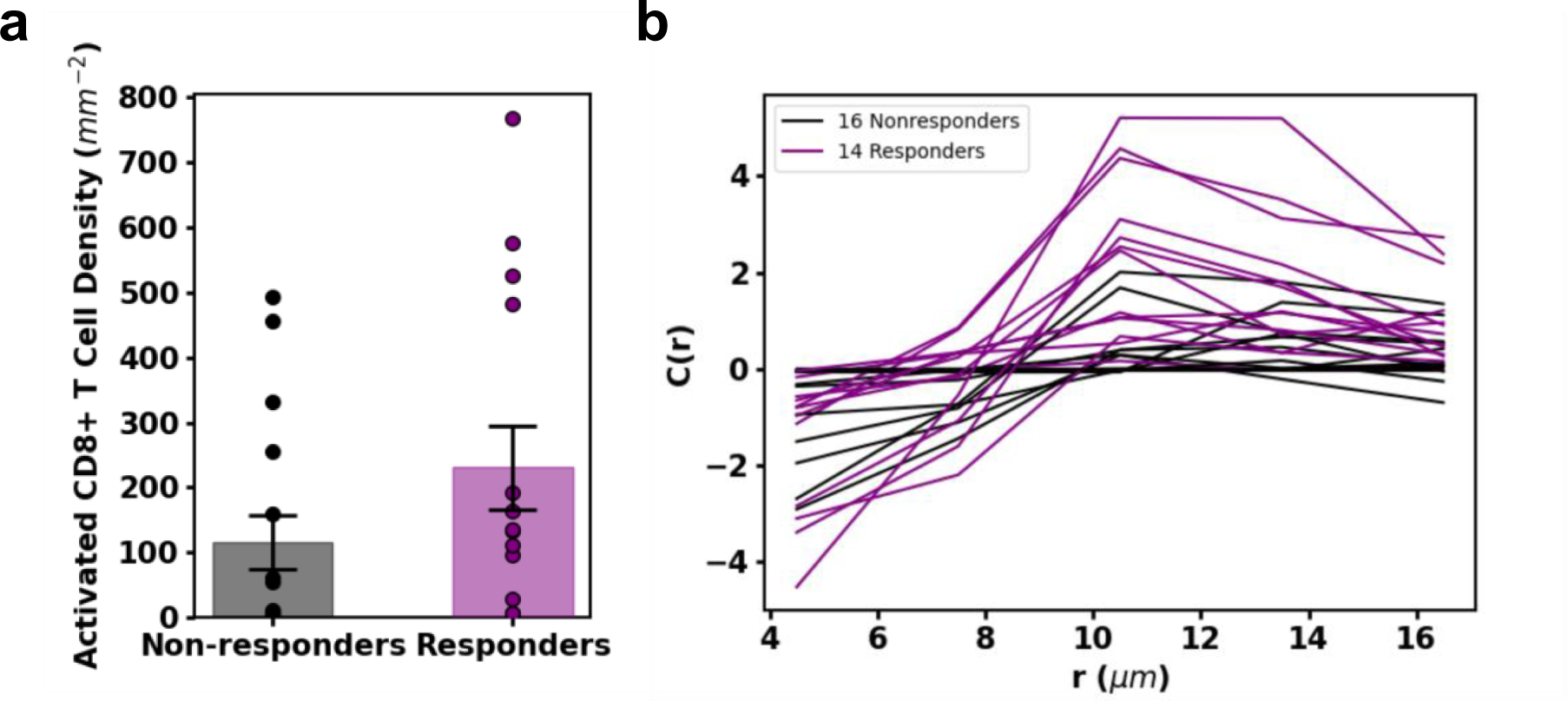
Selection of cell types by analyzing patient slides. **(a)** Activated CD8+ T cell density in slides corresponding to patients who did or did not respond to ICI therapy. The average activated CD8+ T cell density (represented with box) among responders is higher than that of the non-responders. The averages are different with p-value 0.1504. Because activated CD8+ T densities differ between responders and non-responders, we consider activated CD8+ T cells a relevant cell population. **(b)** The plot of spatial correlation between macrophages/monocytes and activated CD8+ T cells shows that at a distance of 10.5 μ*m* (about the distance to a nearest neighbor), macrophages/monocytes find on average more activated CD8+ T cell neighbors in responders than in non-responders. The average spatial correlation values at 10.5 μ*m* are different with a p-value of 0.0005. This separation between responders and non-responders to ICI in the spatial distribution shows the relevance of the cell types, macrophages/monocytes and activated CD8+ T cells, and their spatial distribution.

Next, we evaluated if specific micro-scale organization of certain cell types in the TMA slides significantly separate responders from non-responders. We used pair correlation function (*C*(*r*)), a widely used approach in statistical physics and materials science (25), to evaluate if cells of the same type or different types cluster or avoid each other within a length scale *r* relative to homogeneous and random spatial distribution of the cells in the slide (Fig. S1b). We computed *C*(*r*) for all the cell types and all possible pairs of the cell types which showed large patient-patient variations (Fig. S1c and Fig. 1b). In order to determine spatial patterns involving specific cell types that might distinguish responders to non-responders, we compared values of *C*(*r*) at *r*=10.5μm for all possible pairs of cell types and found that *C*(*r*) for macrophages/monocytes with activated CD8+ T cells differ substantially (p=0.002) between responders and non-responders, whereas, the *C*(*r*) for most of the other pairs of cell types cannot be well separated (p>0.05, Fig. S1c). The only cells other than CD8+ T cells and macrophage/monocytes that showed *C*(*r*) values substantially (p=0.049) different between responders and non-responders are CD4+ T cells and stromal endothelial CD31+ cells. Stromal and immune cell interactions can influence tumor growth and metastasis via cytokines and tumor cell differentiation (26). In this study, we focused on growth and lysis of tumor cells by CD8+ T cells, leaving the influence of stromal and immune cell in tumor growth as a future direction. Based on the above analyses we reasoned that melanoma cells, activated CD8+ T cells and macrophage/monocytes and mechanisms involving interactions between these cell types give rise to the differences in tumor growth in the patients who went through the ICI therapy. In order to evaluate the roles of different mechanisms with which these cells can interact to regulate tumor growth we developed a spatially resolved mechanistic ICS model as we describe below.

#### Development of spatially resolved interacting cell system (ICS) model

We developed a spatially resolved model to describe time evolution of melanoma cells and specific immune cells we identified from our data analysis. The model is set up on a 1×1 mm^2^ two-dimensional simulation box divided into smaller *l*_0_ × *l*_0_ (=10 × 10 μm^2^) chambers. We considered activated CD8+ T cells, exhausted CD8+ T cells, tumor associated macrophages (TAMs) and melanoma cells (Fig. 2a) where these cells interact and move spatially with specific rules, and proliferate, die, or differentiate. The rules are based on experimental observations reported in the literature or on previous computational modeling efforts. We briefly describe the rules used in the model below and provide biological justifications for the rules, and parameter values associated with the rules in Table 1 and in the Materials and Methods section.

**Figure 2.**
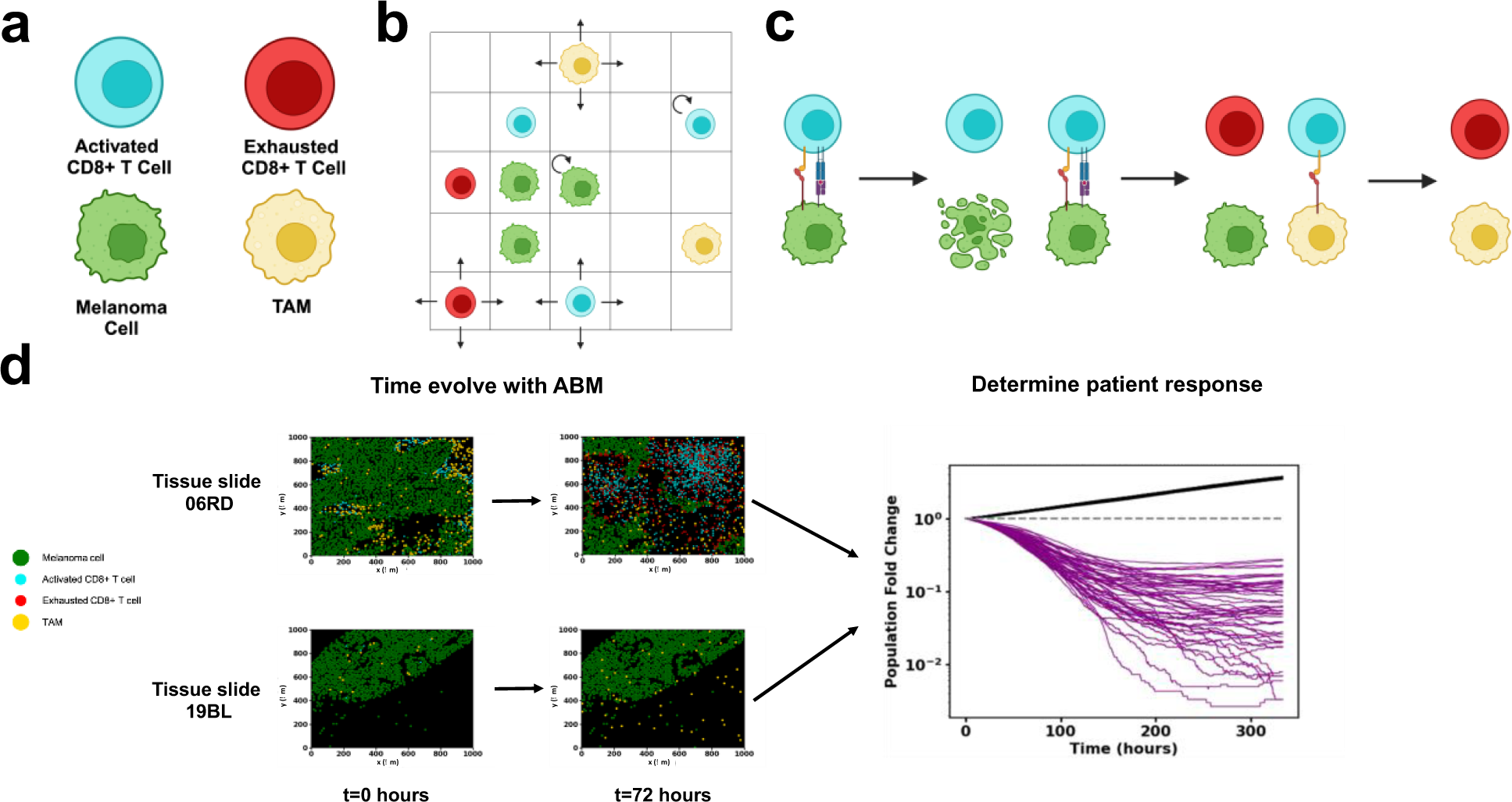
Schematic of model and approach to parameter estimation and hypothesis testing. **(a)** The 4 cell types in the model: including melanoma cells, activated CD8+ T cells, exhausted CD8+ T cells and TAMs. **(b)** Schematic depicting melanoma cells, macrophages, activated CD8+ T cells and exhausted CD8+ T cells on the ICS lattice. CD8+ T cells and macrophages are free to diffuse around the slide. **(c)** Cell-cell interactions in the model include lysis of melanoma cells by activated CD8+ T cells at rate *l*, exhaustion of activated CD8+ T cells by melanoma cells at rate *b*_*c*_ and exhaustion of activated CD8+ T cells by TAMs at rate *b*_*M*_. Rates and rules can be found in Table 1. Figures (a), (b) and (c) created with BioRender.com. **(d)** Plots of positions of melanoma cells, macrophages, cytotoxic CD8+ T cells, and exhausted CD8+ T cells from IMC data of human melanoma biopsies taken before ICI treatment and corresponding computationally time-evolved samples after two days. (Top) IMC slide 06RD corresponding to a responder. (Bottom) IMC 19BL corresponding to a non-responder. (Right) Melanoma cell population trajectories in time for fifty samples for each of two slides. The 19BL samples display cancer growth showing non-response and the 06RD samples display cancer regression which is an example of response.

**Table 1:**
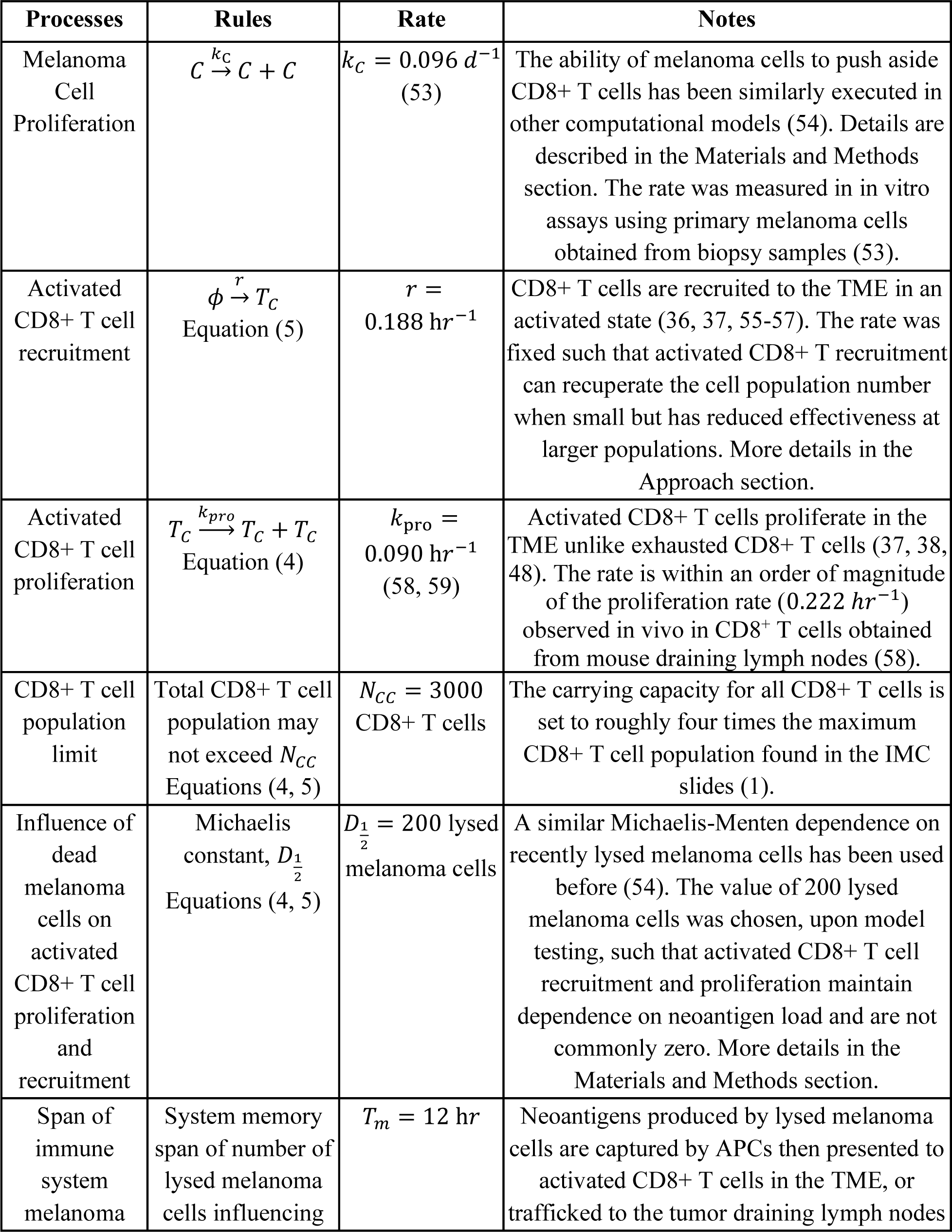
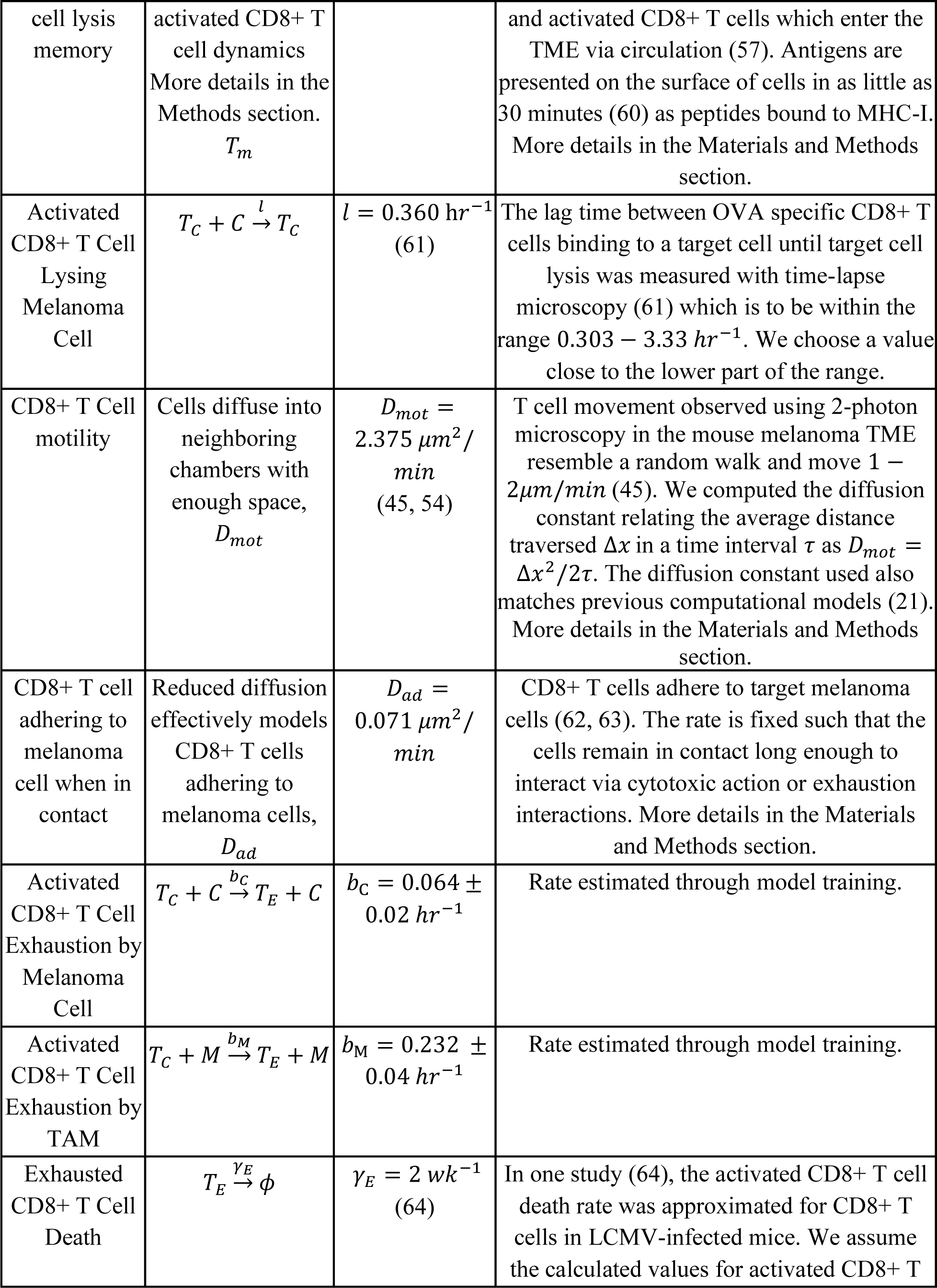
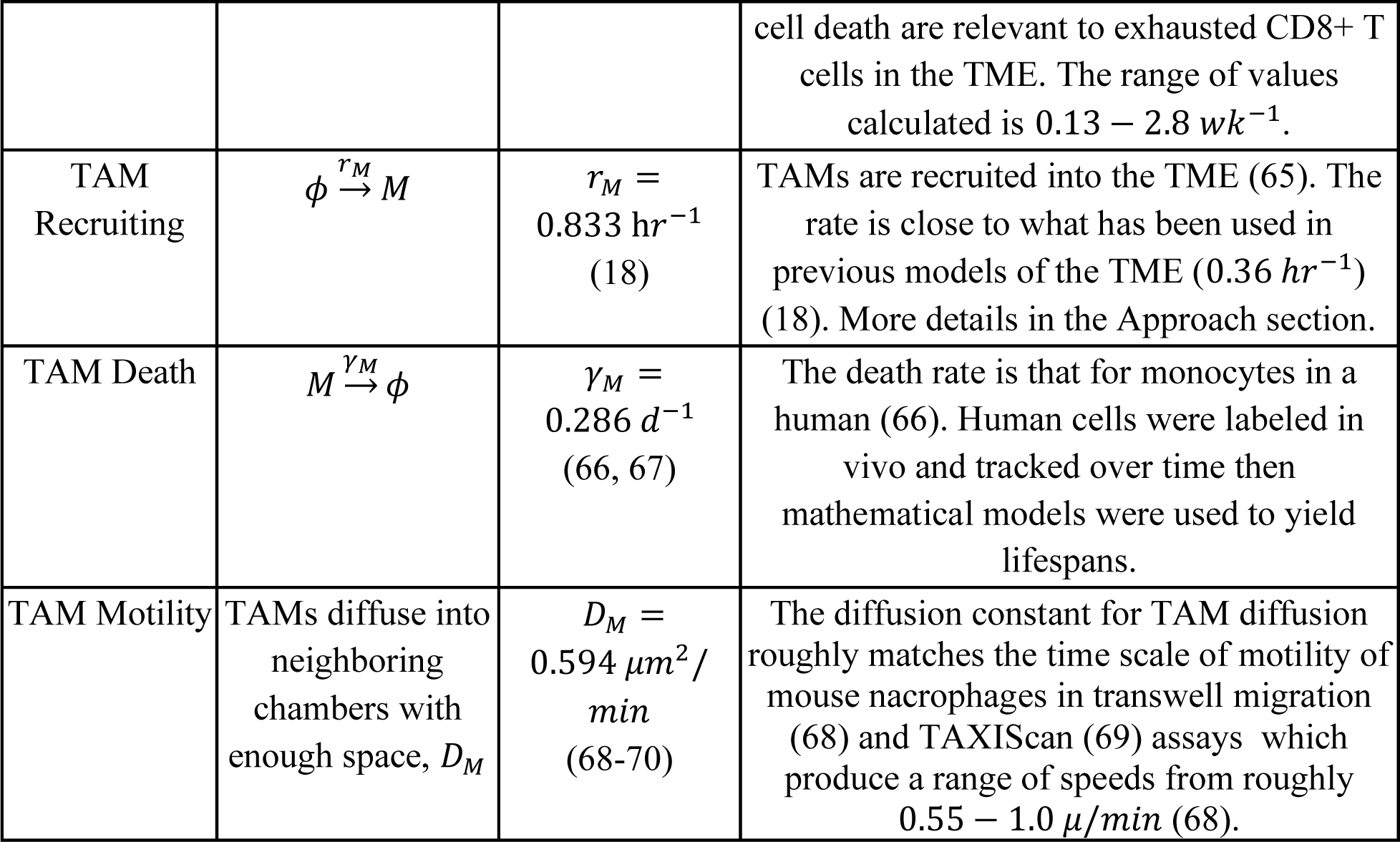
Processes, rules, and parameters used in the ICS model. All parameters are held fixed through all simulations except for the two rates which are estimated through the training process: the exhaustion rates of activated CD8+ T cells by both melanoma cells (*b*_*c*_) and TAMs (*b*_*M*_).

In the model, all the four cell types can occupy chambers in the simulation box obeying local occupation limits due to the cells’ physical sizes. Melanoma cells proliferate (Fig 2b) and are lysed by activated CD8+ T cells when the melanoma cells are in contact with the CD8+ T cells in the same or neighboring chambers (Fig. 2c). When in contact with melanoma cells, activated CD8+ T cells can become exhausted. Activated CD8+ T cells proliferate creating new activated CD8+ T cells in the same chamber. Activated CD8+ T cells are also recruited into a randomly chosen chamber. The rates of proliferation and recruitment of activated CD8+ T cells depend on the number of melanoma cells lysed within a time interval. The exhausted CD8+ T cells do not lyse melanoma cells or proliferate, and die. TAMs, similar to melanoma cells, induce CD8+ T cell exhaustion when those are in contact with activated CD8+ T cells in the same chamber. Spatial motility of CD8+ T cell and TAM are modeled as diffusive hops to the nearest neighboring chambers. More details are given in the Materials and Methods section and Table 1.

#### Modeling the time evolution

We approximate the randomness in cell movements, cell proliferation, cell death, and cell differentiation events as Markov processes where the current state only depends on the previous state of the system for simplicity. The time evolution is performed by a kinetic Monte Carlo simulation approach (details in Materials and Methods).

#### Model Training

The time evolution of our model depends on the values of the model parameters. The order of magnitudes of many of the parameters are known from previous experiments and modeling efforts, however, some of the parameters that substantially affect the tumor growth such as the rate of exhaustion of activated CD8+ T cells by melanoma cells or macrophages are not known. We estimated these parameters using the patient response outcomes in the following way. The initial locations and numbers of the melanoma and immune cells in our model are obtained from the TMA slides, and the locations and numbers of these cells change as the model is evolved over time. We computed the fold change of the total number of melanoma cells from the initial configuration at t=0 with that of at an end time T, where a fold change of less (or greater) than 1 designates the patient associated with the TMA slide as a responder (or non-responder) (Fig. 2d). Since the time evolution is stochastic, the above fold change can vary across multiple simulations of the same initial condition with the same model parameters. We therefore define a frequency variable that quantifies the fraction (*f_i_* (θ)) of the simulations of a model, defined by parameter set *θ,* from the same initial condition from a slide *i* agrees with the clinical outcome for the corresponding patient. We then calculate an appropriate prediction success score for this model with parameter set *θ* to quantify the model’s success across all slides. Finally, maximizing this success score yields the data-optimized parameter set which is used throughout the rest of the study. Additional details can be found in the Materials and Methods section.

#### Hypothesis testing

We set up a hypothesis testing framework for evaluating different mechanisms that can potentially underlie the regulation of tumor growth by the immune cells considered in our model using model simulations, IMC datasets, and the clinical outcome data. Alternate mechanisms (e.g., TAMs do not exhaust activated CD8+ T cells) can be tested by fixing some of the model parameters in the base model (the model described up to this point) to zero to create test models. To compare these test models to the base model, we must first train the new models as we did the base model (Fig 3a). Once these test models are trained, we then utilize bootstrapping to generate new datasets. Finally, comparing the prediction success scores for each dataset between the base model and a given test model, we determine which model is more successful at predicting response (Figs. 3b, S2). Further details are provided in Materials and Methods section.

**Figure 3.**
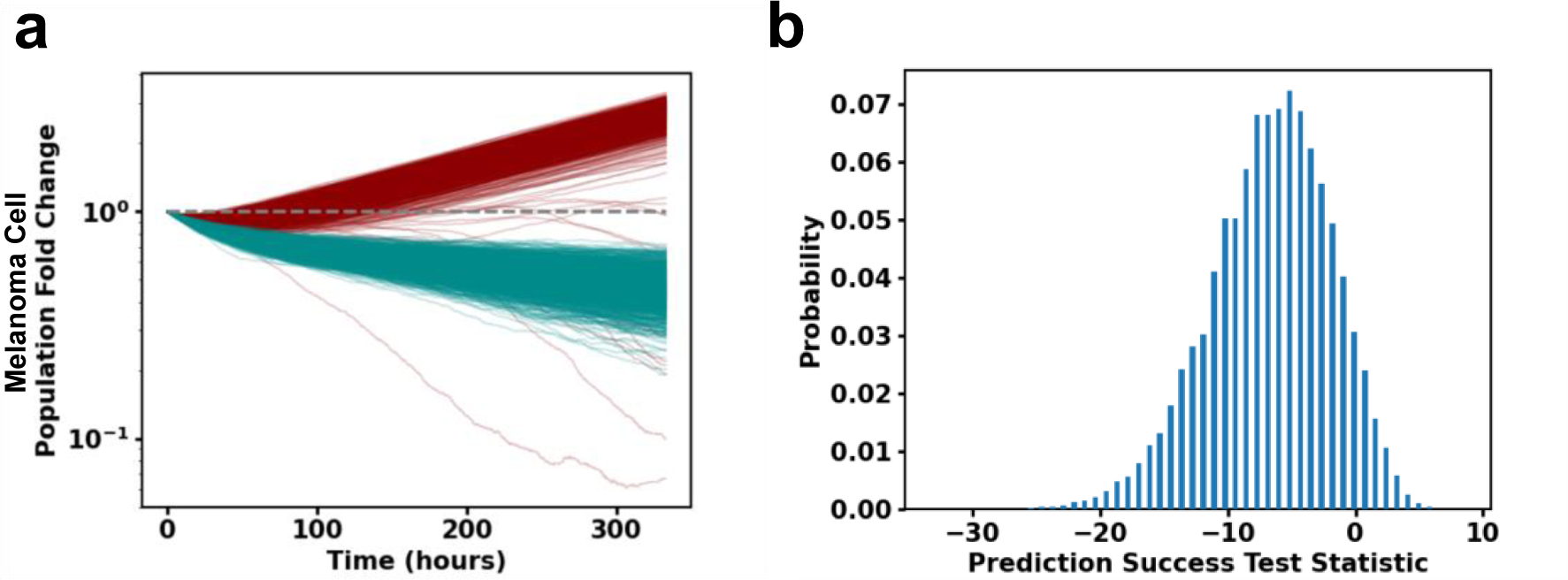
Hypothesis testing with ICS model and patient response data suggests TAM and melanoma cell induced exhaustion of CD8+ T cells regulate response to ICI drugs. **(a)** Melanoma cell fold change (log-linear plot) trajectories plotted as a function of time for 1000 simulations of slide 33RD performed with (dark red) the trained, full model and with (dark cyan) the trained model with TAM exhaustion of activated CD8+ T cells turned off. **(b)** Test statistic distribution with 100,000 bootstraps to test the hypothesis “prediction power of the full model is the same as in the model without TAM exhaustion of activated CD8+ T cells”. Hypothesis is rejected with p-value 0.070. The test statistic distribution for the model without melanoma cell exhaustion can be found in Figure S2.

## Results

### 1. Exhaustion of CD8+ T cells induced by TAMs and melanoma cells regulate response to ICIs

We tested several hypotheses pertaining to the mechanistic interplays between CD8+ T cells, TAMs, and melanoma cells that have been previously explored in experimental investigations with animal models, in vitro and ex-vivo studies by applying our approach to the IMC and the patient response data in (6). To test if macrophages play a pro-tumor role in the TME, we evaluated the hypothesis that exhaustion of CD8+ T cells by TAMs and melanoma cells plays a pro-tumor role and influence the response to ICI drugs in patients. We compared the base model with alternate models (representing hypotheses) where either the rates with which TAMs or melanoma cells exhaust CD8+ T cells when these cells interact is set to zero. The absence of CD8+ T cell exhaustion by either melanoma cells or TAMs in the alternate models led to larger decreases in the fold change of the melanoma cells compared to that in the base model (Fig. 3a). This is due to the presence of more activated CD8+ T cells in the TME which sustained cytotoxicity towards the melanoma cells for the duration of the simulation. We evaluated the consequence of removing exhaustion of CD8+ T cells by melanoma cells and TAMs as in the alternate models on their ability to predict response to ICI drugs within our hypothesis testing framework. We found suggestive p-values of 7% (TAM exhaustion of CD8+ T cells to zero) and 9% (melanoma cell exhaustion of CD8+ T cells to zero) respectively rejecting the alternate hypotheses; therefore, the alternate models are less successful in predicting the patient outcome compared to the base model (Fig. 3b, S2). Thus, this result points to the importance of the exhaustion of CD8+ T cells by TAMs and melanoma cells in regulating tumor growth and response to ICI drugs in melanoma.

### 2. The initial (pre-treatment) spatial organization of immune cells determines melanoma cell growth in the TME

Characterization of the trained ICS dynamics yields insight into how the TME might evolve in vivo. In particular, we find that the initial spatial distribution of activated CD8+ T cells and TAMs in the ICS model impacts the average dynamics over all simulations (trajectory average) of melanoma and activated CD8+ T cells for several patient slides. To evaluate the role of the initial spatial distribution of the immune cells in the TME, we first uniformly and randomly distributed activated CD8+ T cells or TAMs throughout the area of each slide to generate altered initial distributions of the immune and melanoma cells. We then investigated the ICS time-evolutions of the altered initial distributions and their ability to predict the patient responses associated with those slides. We describe two such cases below.

Consider slide 33RD corresponding to a patient which did not respond to ICI therapy. The base model showed a net increase in the average number of the melanoma cells at the final time (about 14 days) and correctly predicted the patient response in 98.6% of 1000 simulations performed of the slide TME (Fig 4a). However, when the initial spatial distribution of the CD8+ T cells was altered by seeding the cells uniformly and randomly throughout the slide, there was a net decrease in the total number of melanoma cells over the duration of the simulation and 100% of the 1000 simulations of the altered initial condition incorrectly predicted the slide to be a responder (Fig. 4a). Inspection of the spatial organization of the melanoma, TAMs, and activated CD8+ T cells in slide 33RD (Fig. 4c), shows that a large portion of the melanoma and the activated CD8+ T cells are spatially segregated which could limit the access of the CD8+ T cell population to the majority of the melanoma cells, whereas, when the activated CD8+ T cells are distributed homogeneously in the altered initial distribution, most of the melanoma cells can be accessed and eliminated by the CD8+ T cells. This leads to a net decay of the melanoma cell population and causes the model to predict the slide to be a responder.

**Figure 4.**
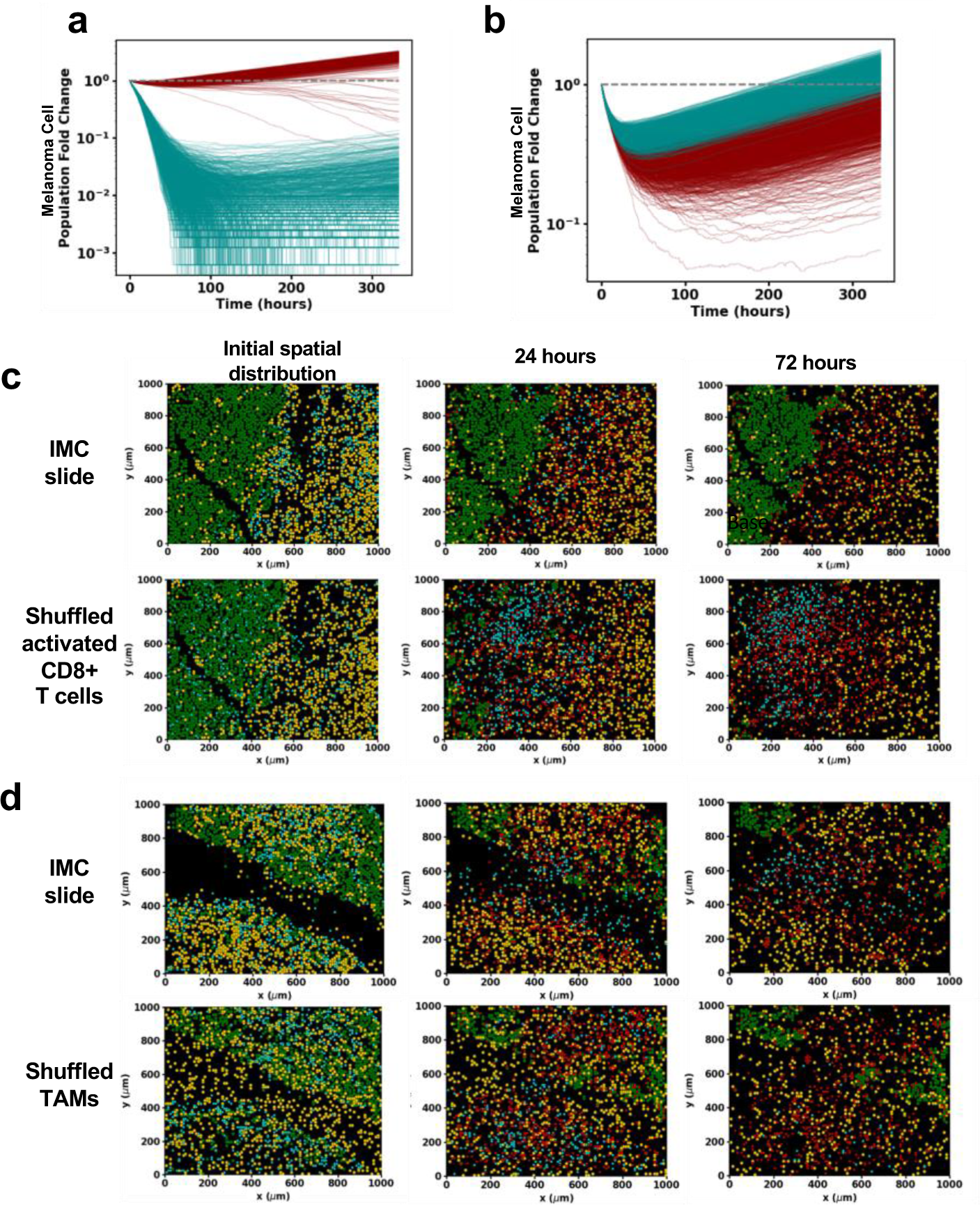
The pre-treatment spatial organization of activated CD8+ T cells, TAMs, and melanoma cells determine tumor cell dynamics in the ICS model. **(a)** Melanoma cell fold change as a function of time on a log-linear plot for 1000 samples of slide 33RD. Contrast the curves for the observed ICI initial condition in dark cyan with those in dark red for the same initial condition but with the initial activated CD8+ T cell positions randomly shuffled. With the rearrangement of the activated CD8+ T cells, the model prediction flips from non-response to response in almost all samples. **(b)** Similar comparison of melanoma cell fold change for 1000 samples of slide 21RD as a function of time; the results for the observed ICI initial condition are in dark red and the results with only the initial TAM positions are randomly shuffled are in dark cyan. The shuffling of the initial TAM positions changes the behavior from responder to non-responder for almost all samples. **(c)** Snapshots of a time evolution of slide 33RD, considered in (a), for one sample using the ICS model (top) and for the same initial condition but with the activated CD8+ T cell positions randomly shuffled (bottom). When the activated CD8+ T cell population is shuffled, it is distributed throughout the melanoma bulk and is thus able to rapidly expand due to its early lysing of melanoma cells. **(d)** Snapshots from a single time evolution of slide 21RD considered in (b) using the ICI initial condition (top) and of the same initial condition but with the TAM spatial distribution randomly shuffled (bottom). The positions of the shuffled TAM cells can more effectively inhibit the expansion of the activated CD8+ T cell population leading to a different outcome. Results for figures 4(a)-(d) were obtained with the trained full ICS model.

In another slide, 21RD, associated with a responder, the time evolution of the initial distribution of the melanoma and immune cells using our ICS model correctly predicts the outcome 98% of the 1000 simulations (Fig 4b). However, when the spatial distribution of the TAMs in slide 21RD is altered by seeding the TAMs randomly and uniformly throughout the slides, only 65 of 1000 simulations show a decrease in the melanoma cell population by final time thus predicting the correct outcome 0.065% times (Fig. 4b). For the slide 21RD, distributing the TAMs homogeneously throughout the slide (Fig 4d) increases mixing between the TAMs and activated CD8+ T cells which increases conversion of exhausted CD8+ T cells and, ultimately, leads to increased growth of the melanoma cell populations and incorrect response prediction for the patient. Clearly, ICI response associated with a slide is dependent on specific features of the initial spatial distribution of cells in the TME. These results point to potential mechanisms with which the initial spatial distributions of the melanoma and immune cells can regulate the tumor growth in the presence of ICI drugs. We further characterize and quantify such spatial mechanisms in the next two sections.

### 3. Intratumoral CD8+ T cells produce fencing of tumor boundaries with exhausted CD8+ T cells

We studied the spatial configurations of the melanoma and the immune cells in the ICS model starting from each of the slides to identify features that may potentially affect tumor dynamics. We found that the system displays an increased accumulation of exhausted CD8+ T cells in contact with melanoma cells, that we term “fencing”. This aggregation of exhausted CD8+ T cells in contact with melanoma cells occurred for simulations initialized with several IMC slides (Fig. 5a). The phenomenon of “fencing” occurred not only in the snapshot configurations shown but persisted on a time scale comparable to (or larger than) that is set by the melanoma cell proliferation rate. We quantified the aggregation of the exhausted CD8+ T cells in contact with melanoma cells in our simulations. We present the analysis in the context of slide 06RD and show results for additional slides in the Supplementary Material (Fig. S3a,b). We defined a collection of exhausted CD8+ T cells in contact with melanoma cells as a fencing cluster of exhausted CD8+ T cells where at least one exhausted CD8+ T cell in the cluster shares the same chamber or a nearest neighboring chamber with a melanoma cell and all the others are in contact with fellow exhausted CD8+ T cells in the same chamber or the nearest neighboring chambers (Fig. 5d). The shape of the clusters of exhausted CD8+ T cells may not always resemble the canonical shape of a fence. We found that for the simulations initiated with the slide 06RD, for configurations at 72 hours, 25% of the exhausted CD8+ T cells reside in such clusters containing at least three exhausted CD8+ T cells (Fig. 5e). In contrast, when the CD8+ T cells in these configurations were permuted randomly with other immune cells (activated CD8+ T cells and TAMs), only 9% of the randomized exhausted CD8+ T cells contribute to the clusters with three or more cells, implicating roles of the spatial distributions of the melanoma and the immune cells and their kinetics in giving rise to these fencing structures. In our simulations, the fencing structures arise because of the exhaustion of active CD8+ T cells in contact with melanoma cells. The exhausted CD8+ T cells that can no longer the lyse the melanoma cells in the fence hinder access of other activated CD8+ T cells to the tumor, thus potentially reducing lysis. Given the generality of such a mechanism underlying the occurrence of fences of exhausted CD8+ T cells, we reasoned these structures should be seen in images of tumor samples of melanoma.

**Figure 5.**
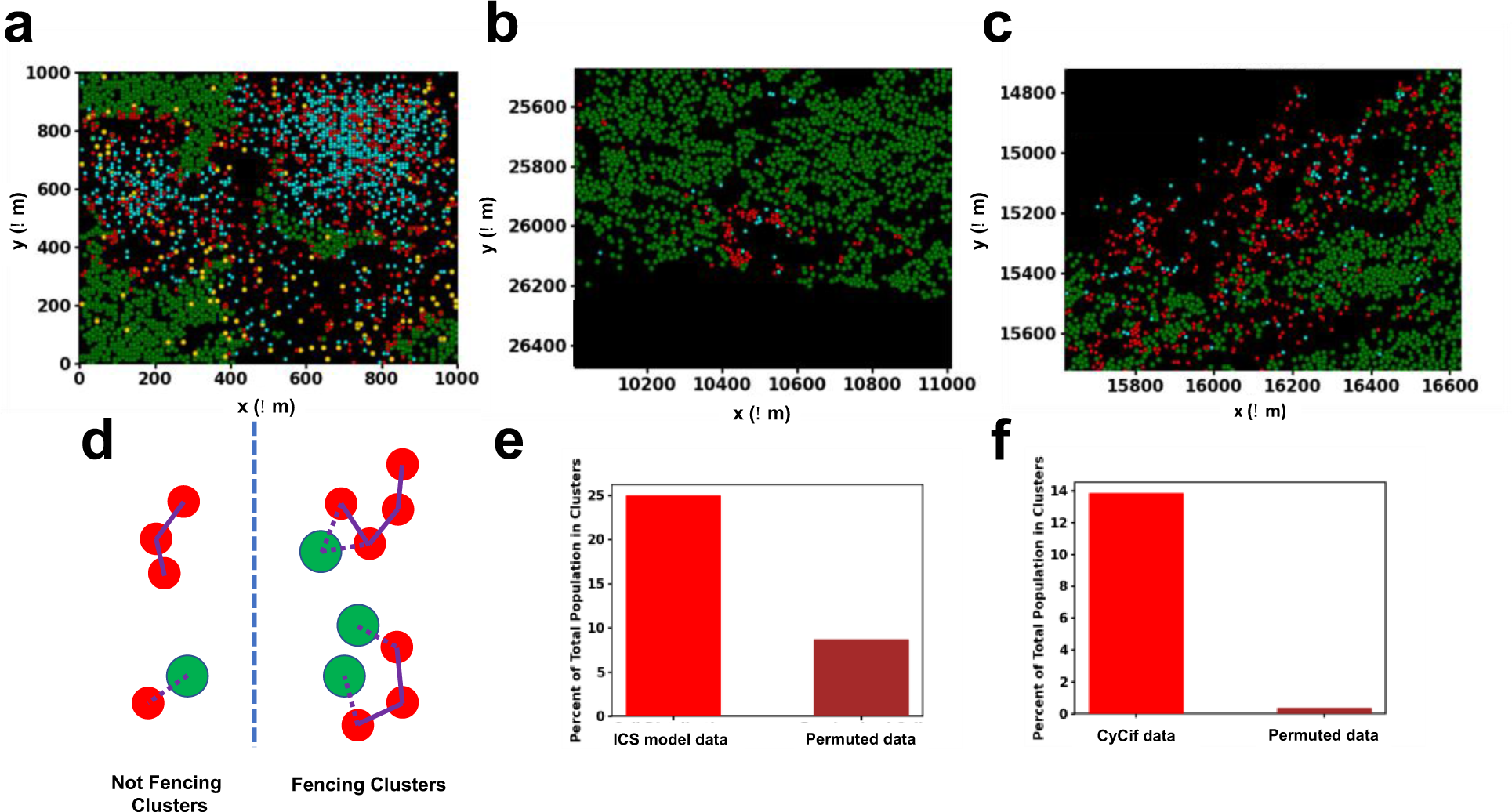
Fencing of tumor boundaries with exhausted CD8+ T cells in the ICS model and CyCIF imaging data of TMA slides from melanoma patients. **(a)** A sample of slide 06RD at 72 hours of simulation shows exhausted CD8+ T cell fencing. **(b), (c)** All melanoma, cytotoxic CD8+ T cells and exhausted CD8+ T cells are plotted for a sub-section of slide MEL06 (1). Exhausted CD8+ T cell fencing in the model is supported by the presence of linings of exhausted CD8+ T cells along the boundary of the tumor in MEL6 blocking access to their cytotoxic counterparts. In general, exhausted CD8+ T cells border the melanoma cells more often than cytotoxic CD8+ T cells. This may lead to a reduction of cytotoxic function in vivo. **(d)** Illustration of forms of fencing and non-fencing clusters with exhausted CD8+ T cells (red) and melanoma cells (green). The two clusters of exhausted CD8+ T cells on the left, considered to be non-fencing, are either not in contact with melanoma cells or do not have enough cells in the cluster. The two clusters on the right meet the criteria for being fencing clusters. **(e)** When the spatial distribution of exhausted CD8+ T cells is randomly permuted with all CD8+ T cells and TAMs in the simulation the percentage of exhausted CD8+ T cells contributing to fencing in slide 06RD at 72 hours is considerably reduced when compared with the actual distribution. **(f)** Percentage of all exhausted CD8+ T cells contributing to fencing in slide MEL06 is similarly greatly reduced when the distribution of exhausted CD8+ T cells is randomly permuted with all identified cells except melanoma cells.

We investigated CyCIF imaging data of melanoma tissue slides obtained from patients at different stages of melanoma progression technique published in a different study by Nirmal et al. (1). The CyCIF imaging data contained over 50 protein markers and identified over 10 different types of immune, stromal, and melanoma cells including melanoma and exhausted CD8+ T cells. We found fencing structures formed by exhausted CD8+ T cells in several regions (two are displayed in Fig. 5b,c). In analyzing the CyCIF data, we took into account the typical physical sizes of the cells in the system, by modifying our definition of nearest neighbors to compute the clusters of exhausted CD8+ T cells in contact with melanoma cells (Fig. 5d). Here, two cells are considered nearest neighbors if their nuclei are within 15 μ*m* of each other. The results of our analysis of slide MEL-6 from ref. (1): 14% of exhausted CD8+ T cells contribute to fencing clusters with three or greater number of member cells; randomly permuting the spatial distribution of the same exhausted CD8+ T cells we found that 0.05% are in the fencing clusters eliminating the possibility that the fencing occurs in random configurations (Fig. 5f). This calculation was also performed with neighbors being defined at other radii and the same qualitative results were found. These results show that exhausted CD8+ T cells preferentially cluster near melanoma cells in vivo, similar to that of observed during the time evolution of the ICS model.

### 4. Growth of melanoma cells in the TME is regulated by stochastic fluctuations

In this section we present results to show how the interplay of the initial spatial distribution of cell types from patient data with stochastic dynamics of the ICS model determines the kinetics of the number of the melanoma cells (Fig. S4a). We will illustrate results from our investigations using two slides as examples, examine their implications, and make several general conclusions. We first examined the kinetics exhibited when the initial condition corresponds to slide 16BL (Fig. 6a,b). The stochastic trajectories describing the fold change of the melanoma cell number over time fluctuate and can be seen to cross at early times (t ≤ τ* approximately 125 hours for 16BL) whereas, as time progresses (e.g., > 125 hours), the overwhelming majority of the 1000 trajectories (except for a few dozen outliers) remain separated (Fig. S4c).

**Figure 6.**
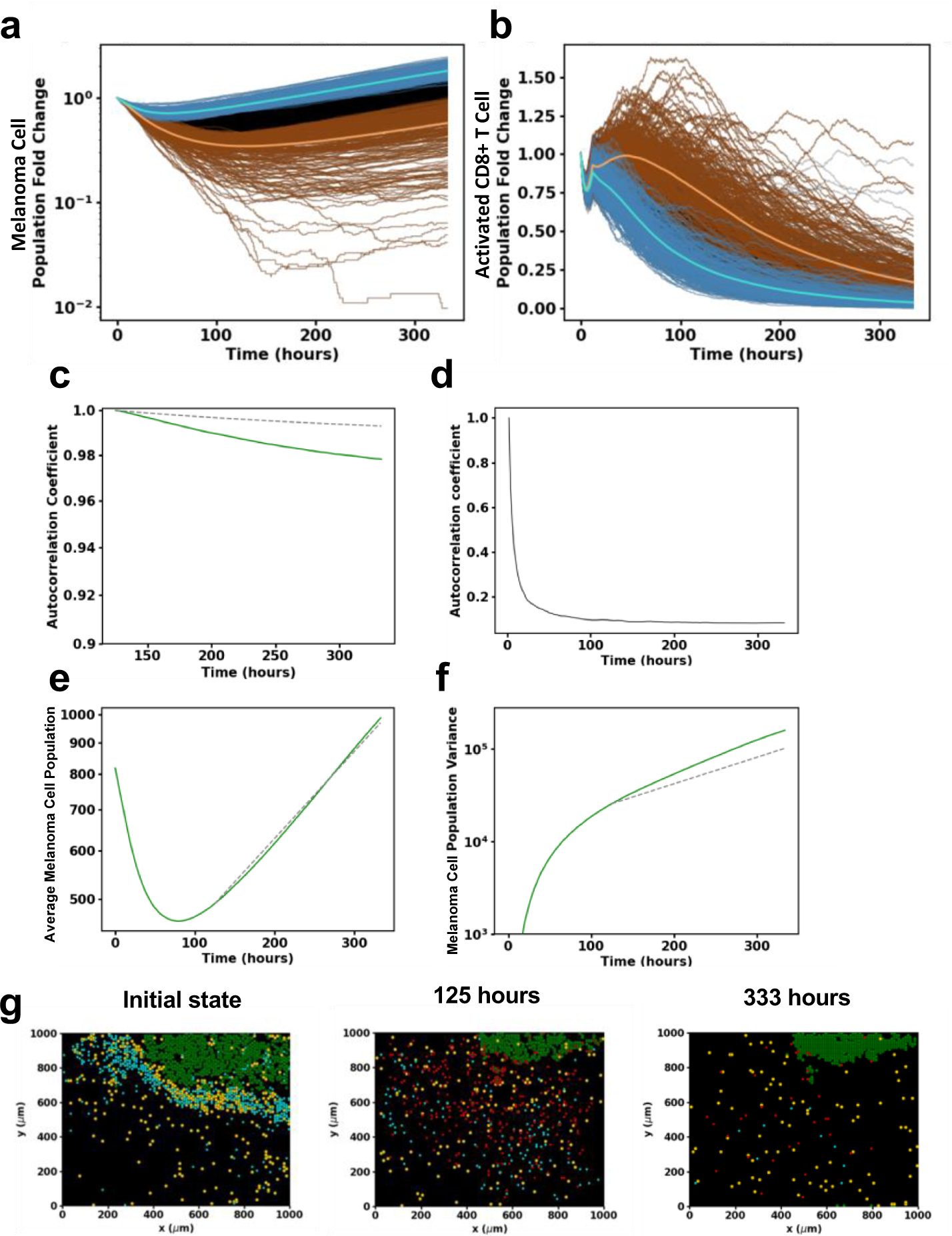
Characterization of the interplay between stochastic fluctuations and initial spatial organization of CD8+ T cells, TAMs, and melanoma cells in regulating tumor growth the ICS model. **(a)** Melanoma cell (log-linear plot) and **(b)** activated CD8+ T cell population (linear-linear plot) trajectories plotted for 1000 samples of slide 16BL. Blue (brown) curves correspond to the top (bottom) 25% melanoma cell populations across all samples at 125 hours. The lighter blue (brown) trajectory corresponds to the average trajectory of the blue (brown) trajectories. The black trajectories represent the other 50% of trajectories which are neither in the blue nor brown subgroups. All samples begin with identical initial conditions set by the patient slide data 16BL. Observe that the samples in the top 25% and bottom 25% of the melanoma population identified at 125 hours remain separated up to the final time. **(c)** The autocorrelation coefficient (Equation 1) of the melanoma cells from 125 hours of the samples plotted with the analytically computed autocorrelation of a simple stochastic birth process Both these autocorrelations decrease very little in the final 175 hours of the simulation. **(d)** The autocorrelation of the ABM from 2 hours to final time. This shows how predictive ability of the future state starting from 2 hours drops off very fast in the early stage. After 125 hours, the autocorrelation drops off very little as in (a). **(e)** A log-linear plot of the mean melanoma cell population as a function of time from the simulations compared with the fitted simple growth process from 125 hours. **(f)** A log-linear plot of the melanoma cell population variance from the simulations compared with the simple growth process variance from 125 hours. The ABM shows good agreement with the simple growth model. The variance is larger for the ABM as there are still some spatial dependences which contribute to variations in population. **(g)** Snapshots of the spatial distribution of the cells from a single time evolution of slide 16BL using the ICS model. The melanoma and activated CD8+ T cells become spatially isolated as the melanoma cell population dynamics transitions to the later growth stage. This isolation persists up to the final time as the activated CD8+ T cells are exhausted, and the melanoma cell population proliferates mimicking a stochastic birth process.

We further characterized the crossing and non-crossing of most of the stochastic trajectories at early and late times respectively using an autocorrelation coefficient function *A*(*t*, *t*_*i*_),

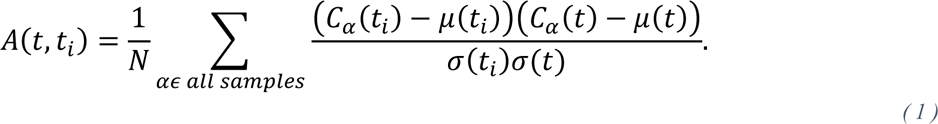

Where *N* is the number of samples, 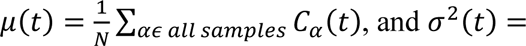 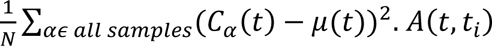 quantifies the correlation between the populations at the current time t and an earlier time *t*_*i*_. The results of the simulations for the slide 16BL are shown in Figure 6c,d; they show a sharp decay of *A*(*t*, *t*_*i*_ = 2*hr*) with t for t > 2 hours capturing the lack of correlation with the initial state or mixing of the stochastic trajectories due to stochastic dynamics whereas, *A*(*t*, *t*_*i*_ = 125*hr*) decays slowly showing the decreased mixing of the stochastic trajectories.

The late-time kinetics of the mean fold change for the trajectories beyond τ* can be well approximated by an exponential growth with a proliferation rate roughly 25% lower than that of the melanoma cell replication rate. The decrease can be attributed to the elimination of the melanoma cells by the remaining small population of activated CD8+ T cells that is still decreasing due primarily to TAM exhaustion. The decreased mixing of the trajectories for t ≥ τ* is reflected in the coefficient of variation (σ/μ, Fig. S4b); it plateaus around a value of 0.4 at late times while the mean melanoma population μ increases exponentially.

The above two observations suggest that the dynamics of the melanoma cell number can be captured by the well-mixed model of a single-variable random birth process, known as the Yule process (27). Fitting the birth rate, the only parameter in the Yule model, to the observed growth rate in the simulations, we compare the analytical expression of the variance of the Yule process with that in the simulations (Fig 6e). The one-variable model variance is somewhat lower than that in the simulations (Fig. 6f). This difference between the ICS model variance and the Yule process can be understood by noting the spatial fluctuations in the number of activated CD8+ T cell population that reduce the effective growth rate. The autocorrelation coefficient function *A*(*t*, *t*_*i*_ = 125*hr*) for the Yule process agrees with that of the ICS model. The fact that we can describe the late stage of the ICS model evolution by a spatially independent model is a striking feature of the model. Next, we examine the qualitative mechanism that leads to the time scale (roughly 125 hours for the slide 16BL) that separates the spatially-stochastic early times and the well-mixed behavior at later time. We reason this is due to a spatial separation between the melanoma and the CD8+ T cell populations. Looking at the spatial configuration at intermediate times (Fig. 7g), we observe that around 125 hours when the average melanoma cell population is near its minimum, the melanoma cells are spatially separated from the activated CD8+ T cell population. The activated CD8+ T cells diffuse through the region with a small probability for encountering and eliminating melanoma cells and only rarely contact the isolated cancer masses. This spatial separation underlies the success of the well-mixed model.

At early times when the CD8+ T cells in closer contact with melanoma cells the lysing of the melanoma cells leads to greater proliferation and recruitment of CD8+ T cells that compensates the exhaustion by TAM and melanoma cells leading to continued decrease of the melanoma cell population. The effectiveness of the feedback mechanism depends on the spatial disposition of the cells in the initial slide and at early times and determines if the spatial separation occurs and if so when. For some of the slides this spatial segregation occurs at early times as in 16BL and similar descriptions are valid (Fig. S6). On the other hand, if the diverse types of cells remain well-mixed as time progresses the feedback persists and there is a decrease in the average number of melanoma cells up to later times and even up to the final time as in slide 06RD (Fig. S5a,b,c) which exhibits mixing as shown by its autocorrelation (Fig S5d). We analyzed stochastic trajectories in other slides which help us make a few general observations described below.

We may break the stochastic dynamics of the other slides into mixing and lower mixing or dispersed trajectory stages similarly to 16BL. The dynamics in all slides except for those with small initial melanoma cell populations may be described by either mixing stages, dispersed stages, or with a transition from the mixing trajectory stage to the dispersed trajectory stage before final time (Fig. S5). For the slides giving rise to stochastic kinetics with the dispersed trajectory stage, the initial activated CD8+ T cell population count and the initial spatial distribution of activated CD8+ T cells determine the time and melanoma cell population count when the slide dynamics transition to the dispersed state. Slides which maintain the mixing stage to final time are characterized by larger activated CD8+ T cell populations at later times.

## Discussion

We integrated cell-level, CyTOF imaging mass cytometry dataset and patient-level response data to ICI treatment in melanoma to develop a mechanistic, spatially resolved ICS model; we used statistical inference to justify and calibrate interactions between melanoma, CD8+ T cells and TAMs in the microscale in the TME. Our simulations allow us to predict responses to ICI treatment and the spatio-temporal development of the TME starting from the snapshot IMC datasets. We used the ICS model to elucidate how specific changes in the spatial co-existence of melanoma and immune cells in the initial state leads to dynamic changes in the spatial organization of the melanoma and the immune cells on microscopic length scales and thus, dramatically affect the growth of the melanoma cell populations. A unique aspect of our model development is the way in which we determined the relevant cell types in our mechanistic ICS model using the IMC and the patient response data, and further estimated model parameters in our model. Furthermore, we studied the time evolution starting from the initial condition provided by the imaging data. This enabled us to identify the important spatial features in much lower dimension (e.g., three cell types) within the large number of cell types, cytokine and chemokines that compose the TME that are crucial in determining the tumor fate.

Inspection of the spatial patterns generated during in our ICS simulations initiated from several TMA slides showed emergence of a fencing pattern of exhausted CD8+ T cells around tumor boundaries. The fencing patterns arise as activated CD8+ T cells encounter melanoma cells and become exhausted. The exhausted CD8+ T cells in the fencing patterns prevent activated CD8+ T cells from invading the interior of the tumor and can potentially protect the tumor cells from CD8+ T cell cytotoxicity. We found confirmatory evidence for such fencing structures formed by exhausted CD8+ T cells (show the biomarkers e.g., TIM3+, Lag3+) in CyCIF imaging data of TMAs obtained from melanoma patients by Nirmal et al (1). This also suggests the underlying mechanism leading to these patterns in our simulations may be operative in melanoma patients. The exhausted CD8+ T cells in the data by Nirmal et al. are likely to arise due to PDL1-PD1 axis induced exhaustion by TAMs and melanoma cells. It will be important to determine if the similar mechanism underlying the fencing pattern of exhausted CD8+ T cells is present in solid tumors other than melanoma, and the differences in the subtypes of CD8+ T cells residing within and outside the fencing structures.

Our investigation found that the spatial segregation of melanoma cells and activated CD8+ T cells plays a pro-tumor role during TME progression in the ICS model. We found that this segregation often occurs at a diminished activated CD8+ T cell population and can mark a transition to a stage which shares characteristics (Fig. S5e), including predictability, with stochastic simple growth. We also observed that spatial mixing of activated CD8+ T cells and TAMs in the microscale tumor tissues increases the probability of the interactions between these cells leading to increased exhaustion of the CD8+ T cells in the TME (Fig. 4c) and subsequent tumor growth. Similar conjectures on the relevance of cellular spatial distributions have been made qualitatively previously (28–34). We were able to deduce such dependencies from the time-evolution of patient slides and our ICS model provided mechanisms that underlie this behavior.

### Limitations of the approach

We ignored several possibly relevant variables in our ICS model for simplicity and due to difficulties in providing validation against experiments. For example, our ICS model did not include the tumor vasculature – the blood vessels network regulating flow of nutrients and waste products in the TME, which is an important factor for tumor growth (35). Our analysis of the IMC dataset determined endothelial cells (CD31+ cells) to be associated with responders to the ICI treatment. Cytokines and chemokines could affect immune cell proliferation, exhaustion, and recruitment in the TME as well as the three-dimensional structure of the tumor that was not explicitly included in the model. Nevertheless, we expect the mechanisms underlying the phenomena we have identified will play a role in the extended models, although quantitative details can differ. These factors could affect the rates of the processes in our model even make them dependent on time. We plan to include these factors supported by relevant, available experimental data in future iterations of this model.

## Materials and Methods

*Analysis of IMC datasets*: These calculations include cell counts, cell densities, and quantifying spatial organization of different cell types with tools such as the two-point correlation.

### Density

The density (σ) of a particular cell type is given by 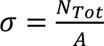. where *N*_*Tot*_ is the total number of cells of that type in the slide and A is the area of the cell tissue in that slide. The area A is calculated by partitioning the slide into a square lattice with lattice constant *a* = 30 μ*m* and then finding all lattice squares which contain at least one cell. These occupied lattice squares comprise the tissue region of the slide and constitute area of the cell tissue.

### Spatial Correlation

We computed the spatial correlation, e.g., between macrophages/monocytes and CD8+ T cells, for a slide in the following way. For each macrophage/monocyte (indexed by i) in the slide, we draw an annular region of radius r and thickness δ (= 3 μ*m*) with the macrophage/monocyte positioned at the center and compute the density of CD8+ T cells in that annular region. Let *n*_*i*(*r*−δ/2,*r*+δ/2)_ be the number of CD8^+^ T cells in the annular region with area A_annulus_=2πrδ; the density σ*_i_*(*r*) of the CD8+ T cells in the annular region surrounding the i^th^macrophage/monocyte is given by

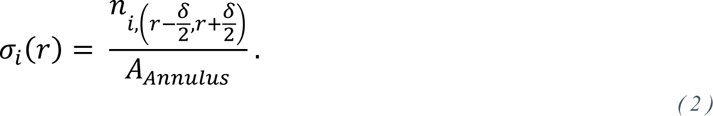

The average density of CD8+ T cells, 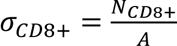 is calculated as discussed earlier. The spatial correlation function is then given by,

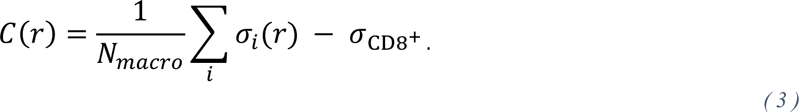

This is then scaled by the average density of CD8+ T cells across all slides σ_cohort,CD8+_. This results in a unitless metric that determines if the clustering of CD8+ T cells around macrophages/monocytes is above or below what is expected from a random distribution and scaled by the average CD8+ T cell density in a slide (Fig. S1b). The scaled correlation function is plotted as a function of r in Figure 2b.

### Model Simulation

The ICS model is a kinetic Monte Carlo simulation implemented on a 100×100 square lattice with spacing 10 μ*m* representing the 1mm×1mm TMAs taken from each patient. We customized SPPARKS Kinetic Monte Carlo Simulator (distributed at spparks.github.io) code to build our model. The references supporting model rules and parameter values are shown in Table 1. The physical extent of the cells constrains the number of cells in each chamber that corresponds to a 10 μ*m* × 10 μ*m* square. CD8+ T cells (active or exhausted) occupy a fourth of a chamber while melanoma cells and macrophages each fill half a chamber. These occupation rules reflect the limits of occupation in the original IMC slides. Two cells are considered in contact if they are in the same chamber or in adjacent chambers that share an edge.

The initial state of the system is obtained from the patient’s TMA by discretizing the cell positions in the image. Over time, cell positions and numbers change: First, melanoma cells proliferate (rate *k*_*c*_) with the location of the daughter depending on the availability of space in progressively larger neighboring regions. The new cell is placed in the same chamber as the cell that is proliferating if space is available, if not then the surrounding 8 chambers, next the adjacent layer of 16 chambers and then the third layer of 24 chambers are checked for available space. When space is not available in a layer, we allow CD8+ T cells in any of the chambers in that layer to be expelled into available space, and check if proliferation can proceed. If no space can be found in the 49-cell neighborhood even by expelling CD8+ T cells, then proliferation does not occur. When multiple chambers are available in each layer one chamber is chosen randomly.

The CD8+ T cells are relocated by similarly iteratively searching their surrounding layers as in melanoma proliferation until a position is found to place them (with no limit). Once open positions for the CD8+ T cells are found, they are placed randomly. This melanoma cell proliferation rule maintains the melanoma cell population’s ability to proliferate even when blocked by a barrier of CD8+ T cells on the tumor periphery.

Activated CD8+ T cells proliferate (36) into the same chamber they occupy if the chamber is not already full. The rate at which activated CD8+ T cells proliferate is dependent on the total population of CD8+ T cells and on the amount of melanoma cells recently lysed:

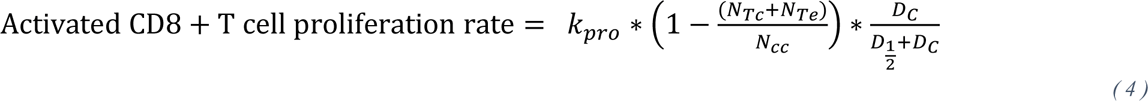

where *k*_*pro*_ is the maximum proliferation rate of activated CD8+ T cells; the carrying capacity N_cc_ limits the maximum value of the sum of the populations of the activated CD8+ T cells (*N*_*Te*_) and exhausted CD8+ T cells (*N*_*Tc*_). The last factor incorporates the dependence of the rate on recently lysed melanoma cells (37, 38). We use a Michaelis-Menten functional form: *D*_*c*_ denotes the number of melanoma cells killed in the previous 12 hours (effective immune system memory span, *T*_*m*_) and 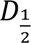 is the Michaelis constant, i.e., when 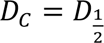, the rate is halved. The larger the melanoma cell death, the greater the probability of neoantigens being created invigorating activated CD8+ T cell proliferation and recruitment by way of APCs. This constitutes positive feedback for activated CD8+ T cell proliferation and recruitment in the TME. APCs are not explicitly modeled by our ICS model but their effects on CD8+ T cell proliferation and recruitment are included.

Activated CD8+ T cells are recruited at a rate with the same dependencies on CD8+ T cell carrying capacity load as the activated CD8+ T cell proliferation rate given by

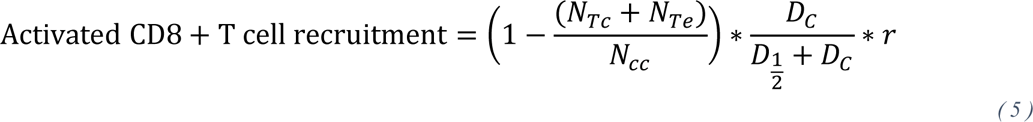

where *r* is the maximum proliferation rate of activated CD8+ T cells. When recruited, activated CD8+ T cells are randomly placed in a chamber with available space for the cell to occupy. TAMs are recruited at a constant rate, *r*_*M*_, and are similarly placed randomly in a lattice chamber with enough space to accommodate the TAM. Exhausted CD8+ T cells and TAMs die at constant rates γ_*E*_ and γ_*M*_ respectively.

In the model, activated CD8+ T cells are transformed into exhausted CD8+ T cells by melanoma cells and TAMs with which they are in contact. In vivo, melanoma cells express PDL1 and PDL2 (31, 39, 40) and chronic PD1 activation in CD8+ T cells leads to exhaustion (41). TAMs express PDL1 and PDL2 in the TME (31) and may directly inhibit activated CD8+ T cells (42–45). In CD8+ T cells, PD1 and CTLA4 expression upregulates upon TCR engagement as well (46, 47). These interactions lead CD8+ T cells to lose cytotoxic and proliferative ability (48).

We account for cell motility in the TMA by allowing activated CD8+ T cells, exhausted CD8+ T cells and TAMs to diffuse on the lattice. A motile cell may move into one of the four nearest neighbor chambers if there is sufficient space for the cell to occupy. The diffusion rate of CD8+ T cells is reduced when in contact with m cells from *D*_*mot*_ to *D*_*ad*_. This is a coarse-grained approximation to the myriad of complex processes that promote contact between CD8+ T and melanoma cells. We apply periodic boundary conditions on our lattice which may be viewed as cell migration to and from other regions of the TME not shown in the data. When crossing a lattice boundary, CD8+ T cells have a 50% chance of being removed from the system.

### Model Training

To train the ICS model, we made use of the fraction of simulations which agree with clinical outcome for model defined by parameter set *θ* and slide *i*, *f_i_* (θ) to build a model prediction success score,

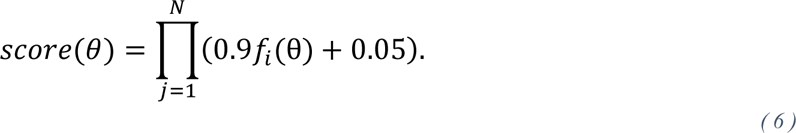

Here, *N* = 30 is the number of patient slides. The vast majority of our patients (approximately 90%) are “compatible” with our ICS model in that our model correctly predicts their drug response with some positive probability. A small number of patients are “incompatible”, and our model will never correctly predict their drug response (e.g., a patient with no activated CD8+ T cells somehow responds to therapy). Because our patient sample is a mixture of compatible and incompatible patients, the success score for the i^th^ slide is a mixture as well. If a patient is compatible then the probability of correct prediction is *f*_*i*_(θ), and if a patient is incompatible then the probability of correct prediction is 1/2 (i.e., no better than random). This leads to the expression shown within the product operator: 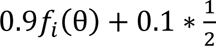. The model which produces a higher success score, fits the patient data more effectively. If we vary model parameters and calculate the score for all parameter sets, we can compare them and find the parameter set which maximizes the prediction success score (49).

By perturbing different model parameters independently, we find that the model is particularly sensitive to variations in the rates at which activated CD8+ T cells are exhausted by melanoma cells and TAMs. Because of their effects on prediction success score, these parameters were chosen to be varied as we optimized the model on the prediction success score. This yields the data-optimized parameter set which is used throughout the rest of the study.

### Hypothesis Testing

Now that we have the ability to train the activated CD8+ T cell exhaustion parameters in our two-parameter ICS response-to-therapy prediction model, we can mimic standard testing procedures commonly used in likelihood theory (50) to compare the larger model to smaller nested models of interest. In particular, we can define a smaller model by fixing one exhaustion parameter to zero and re-training the other. Then, using the success scores from each model, and using a nonparametric bootstrap procedure we can compare the two models (i.e., the larger two-parameter model and the nested smaller model). Rejection of the smaller model occurs with a calculated p-value.

Specifically, the parameters are estimated (as described above) using a cost function that minimizes differences across slides between predicted and observed responses to ICI therapy. Then to perform the test, we compute the difference in negative log success scores (DSS) for the entire sample, where each success score is specific to each model. Because DSS measures the difference in goodness of fit of each model, we use DSS as our test statistic for the bootstrap. Lastly, to make the test computationally feasible, we bootstrap over the two-dimensional vector of slide-specific prediction fractions *f*_*i*_(θ_opt_) found at optimal parameters for each model (as opposed to bootstrapping over each individual slides). From multiple bootstrap resamples, we construct the bootstrap distribution of DSS. If the one-sided (1 − α) × 100%confidence interval for DSS does not contain zero, then we reject the hypothesis of equality at the α × 100% level (i.e., the larger model provides a better fit to the data than the smaller model, and this improved fit cannot be readily explained by chance alone).

### Particle Swarm Optimization

A particle swarm optimization algorithm (51) was utilized to search parameter space to minimize the negative log likelihood of the prediction success score. The set cognitive, social and inertial parameters were 2, 2 and 0.6 respectively (52). The PSO was done with 10 particles searching through 10 sweeps and *f*_*i*_(θ) calculated for each parameter set and slide was done with 50 simulations.

### Cell Segmentation

We segment our cell types following (1). With independent cell locations and marker intensities, we may evaluate each cell and its markers independently to categorize them. With the distribution of marker intensities for a given marker across a cell population, we set a threshold above (below) which a cell is considered positive (negative) in that marker. A cell is determined a melanoma cell if it is Sox10+ and S100a+. If a cell is found to be Sox10-, panCK-, and CD31-then we check if it may be a CD8+ T cell. To be an CD8+ T cell, the cell must also be CD3+ and CD8+. We find activated CD8+ T cells by choosing those CD8+ T cells that are PD1+. Finally, an activated CD8+ T cell is found to be exhausted if it is Tim3+ and Lag3+.

### Data Availability

Data used to generate all figures along with videos of the time evolution of single simulations of slides 06RD and 16BL may be found at DOI: 10.5281/zenodo.10206505. Instructions and code to build and execute ICS model simulations can be found at https://github.com/gdag458/Melanoma_ICS. The patient IMC slides produced by (6) are available at https://doi.org/10.5281/zenodo.5903179. Finally, CyCIF melanoma patient data from ref. (1) are available at https://humantumoratlas.org.

## Acknowledgements

We would like to thank Dr. Rajdeep Kaur Grewal for aid in early SPPARKS development and Dr. Andreas Wieland for his comments and discussion regarding model construction.

## Notes

https://github.com/gdag458/Melanoma_ICS

https://zenodo.org/records/10206506

https://doi.org/10.5281/zenodo.5903179

https://humantumoratlas.org

### Competing Interest Statement

The authors have declared no competing interest.

## References

1. A. J. Nirmal et al., The Spatial Landscape of Progression and Immunoediting in Primary Melanoma at Single-Cell Resolution. Cancer Discovery 12, 1518–1541 (2022).

2. S. J. Turley, V. Cremasco, J. L. Astarita, Immunological hallmarks of stromal cells in the tumour microenvironment. Nature reviews immunology 15, 669–682 (2015).

3. M. M. Steele et al., T cell egress via lymphatic vessels is tuned by antigen encounter and limits tumor control. Nature Immunology 24, 664–675 (2023).

4. P. Sharma, S. Hu-Lieskovan, J. A. Wargo, A. Ribas, Primary, adaptive, and acquired resistance to cancer immunotherapy. Cell 168, 707–723 (2017).

5. A. Young, Z. Quandt, J. A. Bluestone, The balancing act between cancer immunity and autoimmunity in response to immunotherapy. Cancer immunology research 6, 1445–1452 (2018).

6. D. Moldoveanu et al., Spatially mapping the immune landscape of melanoma using imaging mass cytometry. Science Immunology 7, eabi5072 (2022).

7. E. Karimi et al., Single-cell spatial immune landscapes of primary and metastatic brain tumours. Nature 614, 555–563 (2023).

8. J.-R. Lin et al., Highly multiplexed immunofluorescence imaging of human tissues and tumors using t-CyCIF and conventional optical microscopes. elife 7 (2018).

9. S. Berry et al., Analysis of multispectral imaging with the AstroPath platform informs efficacy of PD-1 blockade. Science 372 (2021).

10. A. Li et al., Selective targeting of GARP-LTGFβ axis in the tumor microenvironment augments PD-1 blockade via enhancing CD8+ T cell antitumor immunity. Journal for Immunotherapy of Cancer 10 (2022).

11. D. Hammerl et al., Spatial immunophenotypes predict response to anti-PD1 treatment and capture distinct paths of T cell evasion in triple negative breast cancer. Nature Communications 12, 5668 (2021).

12. J.-R. Lin et al., Multiplexed 3D atlas of state transitions and immune interaction in colorectal cancer. Cell 186, 363–381. e319 (2023).

13. D. Schapiro et al., histoCAT: analysis of cell phenotypes and interactions in multiplex image cytometry data. Nature methods 14, 873–876 (2017).

14. Z. Chen, I. Soifer, H. Hilton, L. Keren, V. Jojic, Modeling multiplexed images with spatial-LDA reveals novel tissue microenvironments. Journal of Computational Biology 27, 1204–1218 (2020).

15. Z. Wu et al., Graph deep learning for the characterization of tumour microenvironments from spatial protein profiles in tissue specimens. Nature Biomedical Engineering, 1–14 (2022).

16. V. Milosevic, Different approaches to Imaging Mass Cytometry data analysis. Bioinformatics Advances 3, vbad046 (2023).

17. C. Gong et al., A computational multiscale agent-based model for simulating spatio-temporal tumour immune response to PD1 and PDL1 inhibition. Journal of the Royal Society Interface 14, 20170320 (2017).

18. C. G. Cess, S. D. Finley, Multi-scale modeling of macrophage—T cell interactions within the tumor microenvironment. PLOS Computational Biology 16, e1008519 (2020).

19. A. Konstorum, A. T. Vella, A. J. Adler, R. C. Laubenbacher, Addressing current challenges in cancer immunotherapy with mathematical and computational modelling. Journal of The Royal Society Interface 14, 20170150 (2017).

20. K.-A. Norton, C. Gong, S. Jamalian, A. S. Popel, Multiscale agent-based and hybrid modeling of the tumor immune microenvironment. Processes 7, 37 (2019).

21. J. N. Kather et al., In Silico Modeling of Immunotherapy and Stroma-Targeting Therapies in Human Colorectal Cancer. Cancer Research 77, 6442–6452 (2017).

22. F. Mpekris et al., Combining microenvironment normalization strategies to improve cancer immunotherapy. Proceedings of the National Academy of Sciences 117, 3728–3737 (2020).

23. L. Hutchinson, O. Grimm, Integrating digital pathology and mathematical modelling to predict spatial biomarker dynamics in cancer immunotherapy. npj Digital Medicine 5, 92 (2022).

24. C. G. Cess, S. D. Finley, Calibrating agent-based models to tumor images using representation learning. PLOS Computational Biology 19, e1011070 (2023).

25. P. M. Chaikin, T. C. Lubensky, Principles of Condensed Matter Physics (Cambridge University Press, Cambridge, 1995), DOI: 10.1017/CBO9780511813467.

26. R. J. Seager, C. Hajal, F. Spill, R. D. Kamm, M. H. Zaman, Dynamic interplay between tumour, stroma and immune system can drive or prevent tumour progression. Converg Sci Phys Oncol 3 (2017).

27. M. A. Pinsky, S. Karlin, “6 - Continuous Time Markov Chains” in An Introduction to Stochastic Modeling (Fourth Edition), M. A. Pinsky, S. Karlin, Eds. (Academic Press, Boston, 2011), 10.1016/B978-0-12-381416-6.00006-X, pp. 277–346.

28. H. Kakavand et al., Tumor PD-L1 expression, immune cell correlates and PD-1+ lymphocytes in sentinel lymph node melanoma metastases. Modern Pathology 28, 1535–1544 (2015).

29. G. Saldanha, K. Flatman, K. W. Teo, M. Bamford, A Novel Numerical Scoring System for Melanoma Tumor-infiltrating Lymphocytes Has Better Prognostic Value Than Standard Scoring. The American Journal of Surgical Pathology 41, 906–914 (2017).

30. C. Keun Park, S. Kyum Kim, Clinicopathological significance of intratumoral and peritumoral lymphocytes and lymphocyte score based on the histologic subtypes of cutaneous melanoma. Oncotarget 8 (2017).

31. J. M. Obeid et al., PD-L1, PD-L2 and PD-1 expression in metastatic melanoma: Correlation with tumor-infiltrating immune cells and clinical outcome. Oncoimmunology 5, e1235107 (2016).

32. S. A. Weiss et al., Immunologic heterogeneity of tumor-infiltrating lymphocyte composition in primary melanoma. Human Pathology 57, 116–125 (2016).

33. C. Fortes et al., Tumor-infiltrating lymphocytes predict cutaneous melanoma survival. Melanoma Research 25, 306–311 (2015).

34. H. Song, Y. Wu, G. Ren, W. Guo, L. Wang, Prognostic factors of oral mucosal melanoma: histopathological analysis in a retrospective cohort of 82 cases. Histopathology 67, 548–556 (2015).

35. J. W. Franses, A. B. Baker, V. C. Chitalia, E. R. Edelman, Stromal endothelial cells directly influence cancer progression. Sci Transl Med 3, 66ra65 (2011).

36. A. C. Huang et al., A single dose of neoadjuvant PD-1 blockade predicts clinical outcomes in resectable melanoma. Nature Medicine 25, 454–461 (2019).

37. S. D. Brown et al., Neo-antigens predicted by tumor genome meta-analysis correlate with increased patient survival. Genome Res 24, 743–750 (2014).

38. T. N. Schumacher, R. D. Schreiber, Neoantigens in cancer immunotherapy. Science 348, 69–74 (2015).

39. M. Mandalà, B. Merelli, D. Massi, PD-L1 in melanoma: facts and myths. Melanoma Manag 3, 187–194 (2016).

40. H. Dong et al., Tumor-associated B7-H1 promotes T-cell apoptosis: A potential mechanism of immune evasion. Nature Medicine 8, 793–800 (2002).

41. B. Bengsch et al., Bioenergetic Insufficiencies Due to Metabolic Alterations Regulated by the Inhibitory Receptor PD-1 Are an Early Driver of CD8+ T Cell Exhaustion. Immunity 45, 358–373 (2016).

42. K. Kersten et al., Spatiotemporal co-dependency between macrophages and exhausted CD8+ T cells in cancer. Cancer Cell 40, 624–638.e629 (2022).

43. I. Kryczek et al., B7-H4 expression identifies a novel suppressive macrophage population in human ovarian carcinoma. Journal of Experimental Medicine 203, 871–881 (2006).

44. W. Zou, J. D. Wolchok, L. Chen, PD-L1 (B7-H1) and PD-1 pathway blockade for cancer therapy: Mechanisms, response biomarkers, and combinations. Science Translational Medicine 8, 328rv324-328rv324 (2016).

45. B. Boldajipour, A. Nelson, M. F. Krummel, Tumor-infiltrating lymphocytes are dynamically desensitized to antigen but are maintained by homeostatic cytokine. JCI Insight 1 (2016).

46. Y. Agata et al., Expression of the PD-1 antigen on the surface of stimulated mouse T and B lymphocytes. International Immunology 8, 765–772 (1996).

47. J. G. Egen, J. P. Allison, Cytotoxic T Lymphocyte Antigen-4 Accumulation in the Immunological Synapse Is Regulated by TCR Signal Strength. Immunity 16, 23–35 (2002).

48. C. U. Blank et al., Defining ‘T cell exhaustion’. Nature Reviews Immunology 19, 665–674 (2019).

49. Anonymous, “M–and Z-Estimators” in Asymptotic Statistics, A. W. v. d. Vaart, Ed. (Cambridge University Press, Cambridge, 1998), DOI: 10.1017/CBO9780511802256.006, pp. 41–84.

50. T. Hastie, R. Tibshirani, J. Friedman, “Model Inference and Averaging” in The Elements of Statistical Learning: Data Mining, Inference, and Prediction, T. Hastie, R. Tibshirani, J. Friedman, Eds. (Springer New York, New York, NY, 2009), 10.1007/978-0-387-84858-7_8, pp. 261–294.

51. J. Kennedy, “Swarm intelligence” in Handbook of nature-inspired and innovative computing: integrating classical models with emerging technologies. (Springer, 2006), pp. 187–219.

52. S. E. Selvan et al., Parameter Estimation in Stochastic Mammogram Model by Heuristic Optimization Techniques. IEEE Transactions on Information Technology in Biomedicine 10, 685–695 (2006).

53. D. A. Rew, G. D. Wilson, Cell production rates in human tissues and tumours and their significance. Part II: clinical data. European Journal of Surgical Oncology (EJSO) 26, 405–417 (2000).

54. C. Gong et al., A computational multiscale agent-based model for simulating spatio-temporal tumour immune response to PD1 and PDL1 inhibition. J R Soc Interface 14 (2017).

55. K. E. Yost, H. Y. Chang, A. T. Satpathy, Recruiting T cells in cancer immunotherapy. Science 372, 130–131 (2021).

56. M. H. Spitzer et al., Systemic Immunity Is Required for Effective Cancer Immunotherapy. Cell 168, 487–502.e415 (2017).

57. I. Mellman, D. S. Chen, T. Powles, S. J. Turley, The cancer-immunity cycle: Indication, genotype, and immunotype. Immunity 56, 2188–2205 (2023).

58. C. Kurts, H. Kosaka, F. R. Carbone, J. F. A. P. Miller, W. R. Heath, Class I–restricted Cross-Presentation of Exogenous Self-Antigens Leads to Deletion of Autoreactive CD8+ T Cells. Journal of Experimental Medicine 186, 239–245 (1997).

59. H. Yoon, T. S. Kim, T. J. Braciale, The Cell Cycle Time of CD8+ T Cells Responding In Vivo Is Controlled by the Type of Antigenic Stimulus. PLOS ONE 5, e15423 (2010).

60. L. C. Eisenlohr, L. Huang, T. N. Golovina, Rethinking peptide supply to MHC class I molecules. Nature Reviews Immunology 7, 403–410 (2007).

61. B. Weigelin et al., Cytotoxic T cells are able to efficiently eliminate cancer cells by additive cytotoxicity. Nature Communications 12, 5217 (2021).

62. H. Raskov, A. Orhan, J. P. Christensen, I. Gögenur, Cytotoxic CD8+ T cells in cancer and cancer immunotherapy. British Journal of Cancer 124, 359–367 (2021).

63. M. L. Dustin, The Immunological Synapse. Cancer Immunology Research 2, 1023–1033 (2014).

64. R. J. De Boer, D. Homann, A. S. Perelson, Different Dynamics of CD4+ and CD8+ T Cell Responses During and After Acute Lymphocytic Choriomeningitis Virus Infection 1. The Journal of Immunology 171, 3928–3935 (2003).

65. T. Hourani et al., Tumor Associated Macrophages: Origin, Recruitment, Phenotypic Diversity, and Targeting. Front Oncol 11, 788365 (2021).

66. J. Cosgrove, L. S. P. Hustin, R. J. de Boer, L. Perié, Hematopoiesis in numbers. Trends Immunol 42, 1100–1112 (2021).

67. J. Lahoz-Beneytez et al., Human neutrophil kinetics: modeling of stable isotope labeling data supports short blood neutrophil half-lives. Blood 127, 3431–3438 (2016).

68. C. Zhao et al., DNA methyltransferase 1 deficiency improves macrophage motility and wound healing by ameliorating cholesterol accumulation. npj Regenerative Medicine 8, 29 (2023).

69. H. Yano et al., Reduction of Real-Time Imaging of M1 Macrophage Chemotaxis toward Damaged Muscle Cells is PI3K-Dependent. Antioxidants (Basel) 7 (2018).

70. E. Au - van den Bos, S. Au - Walbaum, M. Au - Horsthemke, A. C. Au - Bachg, P. J. Au - Hanley, Time-lapse Imaging of Mouse Macrophage Chemotaxis. JoVE doi:10.3791/60750, e60750 (2020).

