## Supplemental Figures for "Spatial organization and stochastic fluctuations of immune cells impact clinical responsiveness to immune checkpoint inhibitors in patients with melanoma"

**a**

| | |
|---|---|
| Melanoma | 0.951 |
| all CD8+ | 0.197 |
| Naïve CD8+ | 0.508 |
| Activated CD8+ | 0.163 |
| All CD4+ | 0.602 |
| Naïve CD4+ | 0.199 |
| Activated CD4+ | 0.914 |
| Treg | 0.4 |
| Macrophage/Monocytes | 0.64 |
| CD31+ | 0.669 |
| B Cells | 0.251 |

**b**

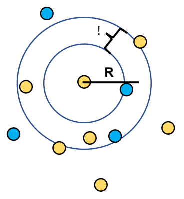


**c**

| | Melanoma Cells | all CD8+ T Cells | Naïve CD8+ T Cells | Activated CD8+ T Cells | All CD4+ T Cells | Naïve CD4+ T Cells | Activated CD4+ T Cells | Treg Cells | Macrophage/Monocytes | CD31+ Cells | B Cells |
|---|---|---|---|---|---|---|---|---|---|---|---|
| Melanoma Cells | 0.567 | 0.641 | 0.634 | 0.565 | 0.551 | 0.186 | 0.915 | 0.273 | 0.363 | 0.57 | 0.206 |
| all CD8+ T Cells | 0.061 | 0.227 | 0.855 | 0.169 | 0.736 | 0.41 | 0.877 | 0.531 | 0.042 | 0.638 | 0.313 |
| Naïve CD8+ T Cells | 0.275 | 0.766 | 0.394 | 0.928 | 0.628 | 0.49 | 0.776 | 0.501 | 0.466 | 0.45 | 0.347 |
| Activated CD8+ T Cells | 0.149 | 0.192 | 0.611 | 0.186 | 0.543 | 0.599 | 0.487 | 0.572 | 0.016 | 0.398 | 0.321 |
| All CD4+ T Cells | 0.373 | 0.908 | 0.374 | 0.876 | 0.728 | 0.385 | 0.865 | 0.315 | 0.235 | 0.049 | 0.369 |
| Naïve CD4+ T Cells | 0.412 | 0.134 | 0.634 | 0.191 | 0.527 | 0.133 | 0.504 | 0.238 | 0.819 | 0.202 | 0.343 |
| Activated CD4+ T Cells | 0.469 | 0.906 | 0.369 | 0.898 | 0.777 | 0.387 | 0.903 | 0.38 | 0.237 | 0.25 | 0.378 |
| Treg Cells | 0.29 | 0.445 | 0.108 | 0.25 | 0.367 | 0.184 | 0.3 | 0.658 | 0.378 | 0.574 | 0.199 |
| Macrophage/Monocytes | 0.106 | 0.004 | 0.96 | 0.002 | 0.061 | 0.434 | 0.083 | 0.529 | 0.026 | 0.224 | 0.216 |
| CD31+ Cells | 0.744 | 0.369 | 0.811 | 0.327 | 0.259 | 0.798 | 0.222 | 0.674 | 0.158 | 0.713 | 0.192 |
| B Cells | 0.154 | 0.176 | 0.97 | 0.171 | 0.72 | 0.693 | 0.741 | 0.344 | 0.241 | 0.406 | 1 |

**Figure S1. Significant variations of spatial relationships between responders and non-responders are yielded from the data. (a)** Table of p-values corresponding to the average density of each species varying between responder slides and non-responder slides. The average activated CD8+ T cell densities are separated with the most confidence with a p-value of 0.16. Responder slides have more activated CD8+ T cells on average. **(b)** Graphic aid for the spatial correlation calculation. The density of blue cells around each yellow cell at radius R in an annulus of thickness $\delta$ is calculated for each yellow cell and averaged across all yellow cells. The total slide density of blue cells is then subtracted from that average to find if the density of blue cells around the average yellow cell is above or below that expected from a random distribution of blue cells. Finally, this value is divided by the average number of blue cells across all slides. **(c)** Table of p-values corresponding to the average spatial correlations at 10.5 $\mu m$ varying between responder and non-responder slides for all permutations of cell type. Values below 0.05 are starred.

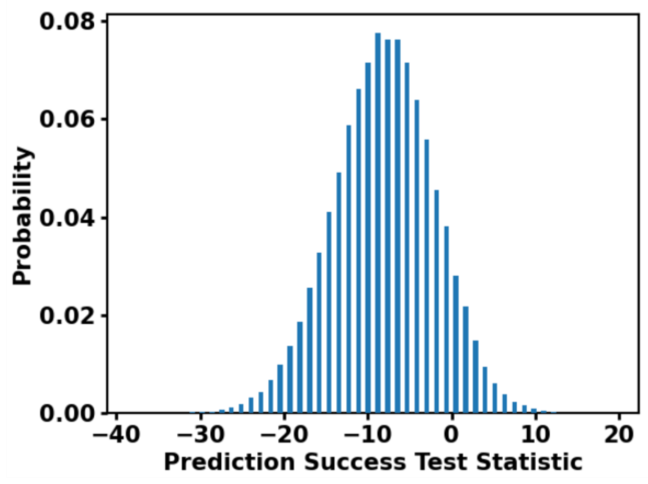


**Figure S2. Melanoma cell exhaustion of activated CD8+ T cell hypothesis test points to relevance of activated CD8+ T cell exhaustion by melanoma cells.** Test statistic distributions with 100,000 bootstraps for hypothesis "prediction power of the full model is the same as in the model without melanoma exhaustion of activated CD8+ T cells". Hypothesis is rejected with p-value 0.088.

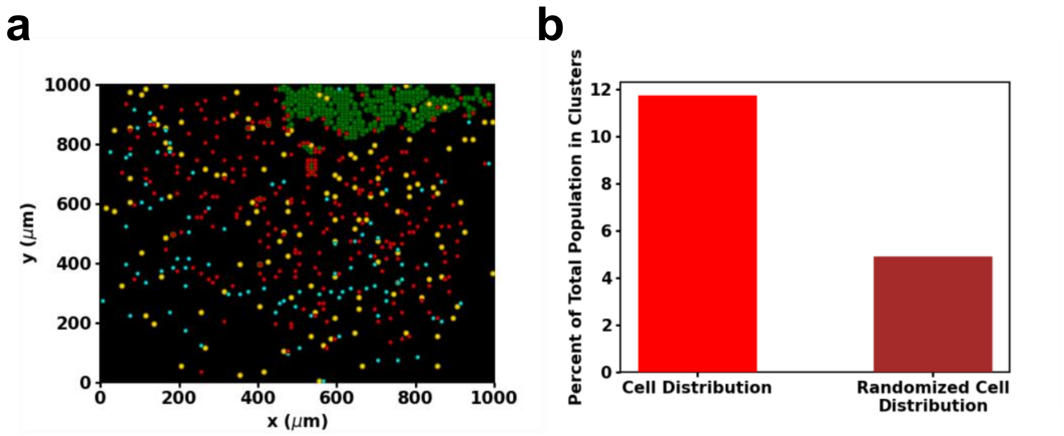


**Figure S3. Exhausted CD8+ T cell fencing exhibited in slide 16BL time-evolution. (a)** Snapshot of the simulated TME of 16BL at 144 hours exhibiting fencing clusters. **(b)** The percentage of exhausted CD8+ T cells in fencing clusters at 144 hours in a simulation of slide 16BL compared to that expected from a randomly permuted distribution of exhausted CD8+ T cells.

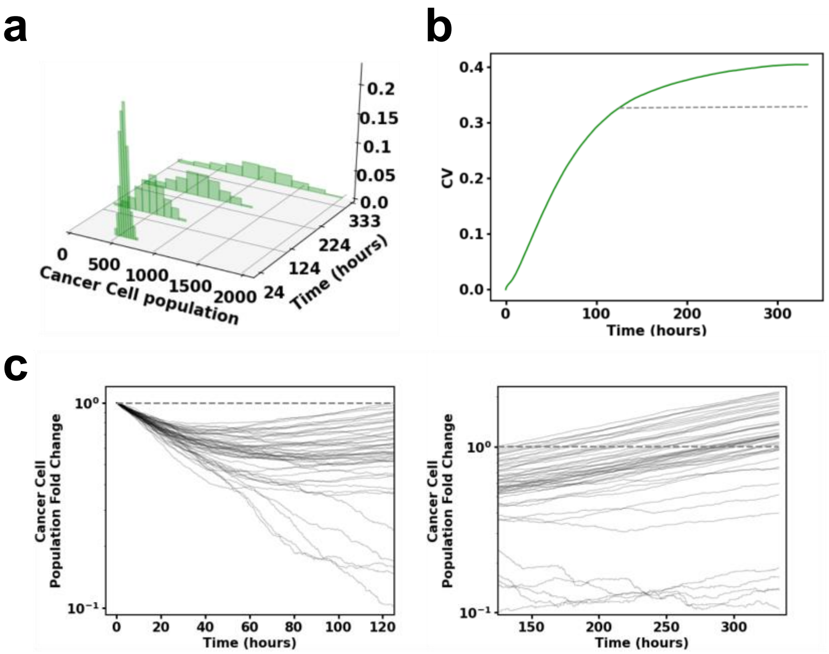


**Figure S4. Characterizing stochasticity in melanoma cell population trajectories corresponding to slide 16BL further. (a)** Cancer cell population distribution yielded by the same 1000 samples as in (a) at increasing times showing how the cancer cell population distribution spreads over time. **(b)** The coefficient of variation (σ/μ) for 1000 simulations of slide 16BL (green) with the fitted Yule coefficient of variation (grey dashed line) from 125 hours. **(c)** 50 cancer fold change trajectories from simulations of 16BL with the trained model plotted from initial time to 125 hours (left) and then from 125 hours to final time (right). We see that early on (around 24 hours to 80 hours), trajectories intersect often whereas at later times (past 125 hours) the trajectories largely remain separated.

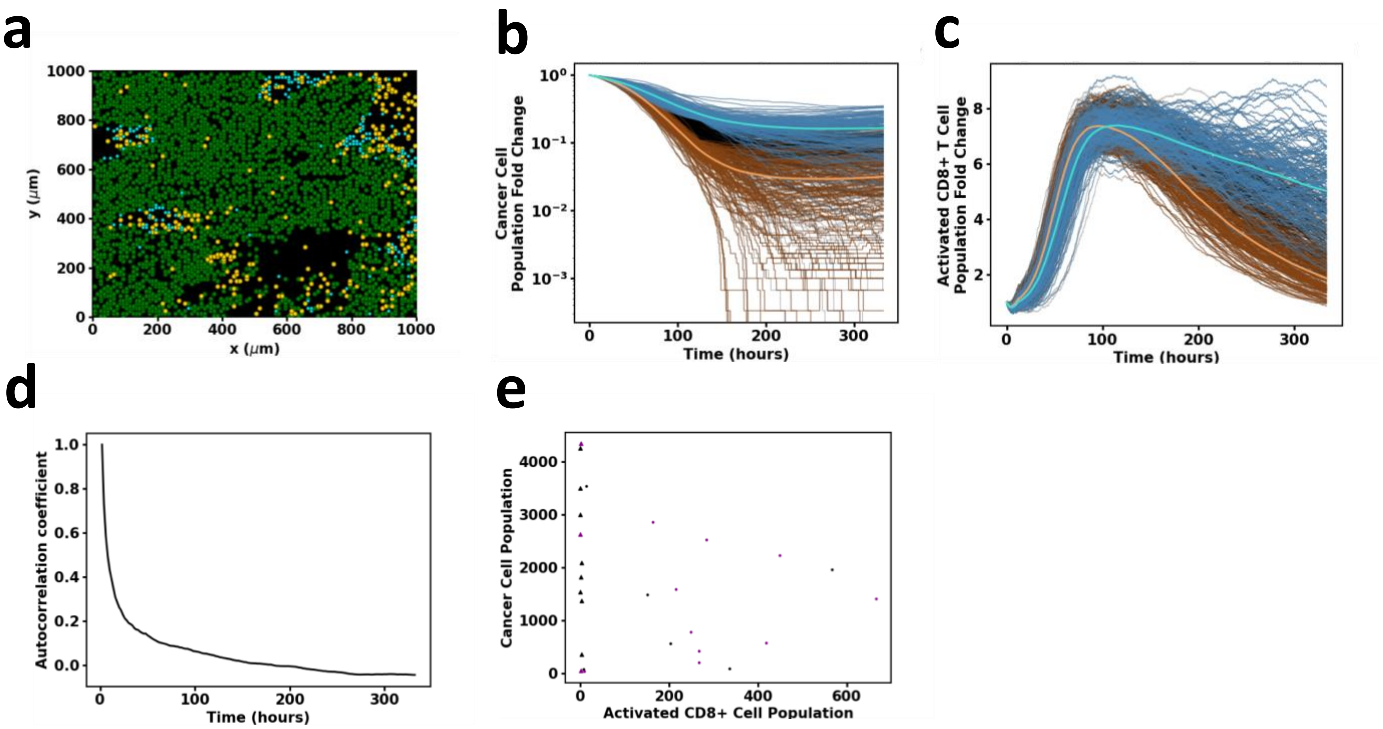


**Figure S5. Stochasticity in ICS-simulated slide 06RD displays the trajectory-mixing stage. (a)** Initial cell spatial distribution for slide 06RD. **(b)** Melanoma cell (log-linear plot) and **(c)** activated CD8+ T cell population (linear-linear plot) trajectories plotted for 1000 samples of slide 06RD. Blue (brown) trajectories are those which compose the top (bottom) 25% melanoma cell populations across all samples at 125 hours. The lighter blue (brown) trajectory corresponds to the average trajectory of all of the blue (brown) trajectories. The black trajectories represent the other 50% of samples. All samples begin with identical initial conditions set by the patient slide data 06RD. Notice how sample trajectories mix until late times unlike in simulations of slide 16BL at late times. 06RD kinetics are also characterized by maintenance of a larger activated CD8+ T cell population until late times. **(d)** The autocorrelation of cancer cell population from 2 hours to final time for slide 06RD. The autocorrelation quickly drops showing that there is low predictive power of future state given initial cancer cell population for this slide. **(e)** Average cancer cell population across 300 samples for every slide at 25 hours plotted against the average activated CD8+ T cell population over the same samples at 25 hours. Purple (black) markers delineate responders (non-responders) and triangles (crosses) mark slides which have (not) transitioned into the simple growth stage by 25 hours. Those slides which start as simple growth processes can be separated out by average activated CD8+ T cell population.

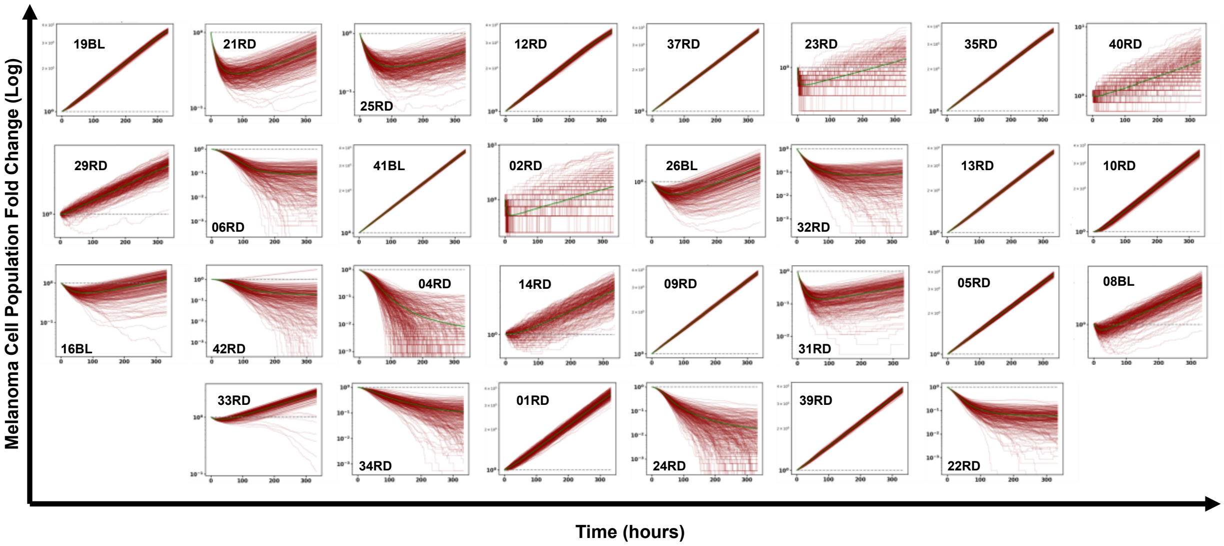


**Figure S6. All slide melanoma cell population trajectories.** Log-linear plots of cancer cell population trajectories for 300 samples simulated initialized by each IMC slide. The average trajectory for each slide is plotted in green.
